# An engineered prodrug selectively suppresses β-lactam resistant bacteria in a mixed microbial setting

**DOI:** 10.1101/2024.08.02.606422

**Authors:** Addison M. Duda, Helena R. Ma, César A. Villalobos, Sophia A. Kuhn, Katherine He, Sarah R. Seay, Abigail C. Jackson, Christine M. Suh, Elena A. Puccio, Deverick J. Anderson, Vance G. Fowler, Lingchong You, Katherine J. Franz

## Abstract

The rise of β-lactam resistance necessitates new strategies to combat bacterial infections. We purposefully engineered the β-lactam prodrug AcephPT to exploit β-lactamase activity to selectively suppress resistant bacteria producing extended-spectrum-β-lactamases (ESBLs). Selective targeting of resistant bacteria requires avoiding interaction with penicillin-binding proteins, the conventional targets of β-lactam antibiotics, while maintaining recognition by ESBLs to activate AcephPT only in resistant cells. Computational approaches provide a rationale for structural modifications to the prodrug to achieve this biased activity. We show AcephPT selectively suppresses gram-negative ESBL-producing bacteria in clonal populations and in mixed microbial cultures, with effective selectivity for both lab strains and clinical isolates expressing ESBLs. Time-course NMR experiments confirm hydrolytic activation of AcephPT exclusively by ESBL-producing bacteria. In mixed microbial cultures, AcephPT suppresses proliferation of ESBL-producing strains while sustaining growth of β-lactamase-non-producing bacteria, highlighting its potential to combat β-lactam resistance while promoting antimicrobial stewardship.

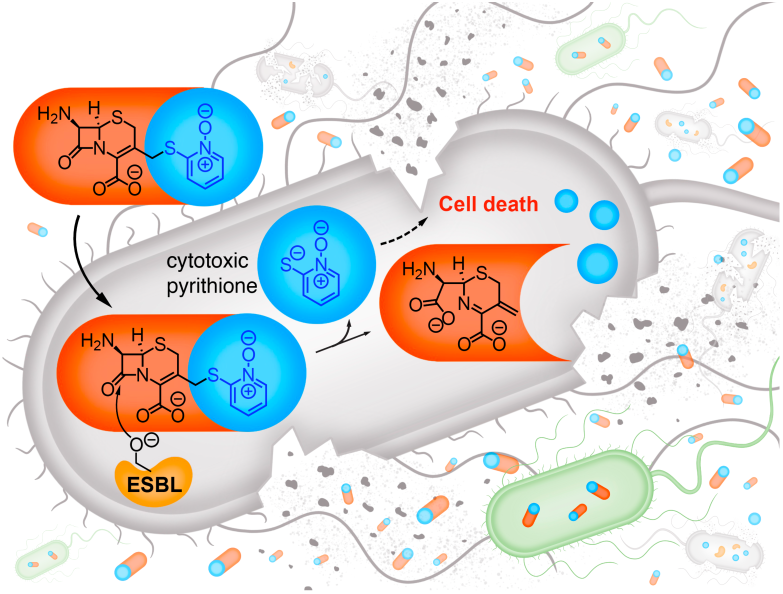

## Introduction

Drugs containing a β-lactam core are the most prescribed antibiotics in the United States.^1^ Resistance to β-lactam antibiotics, however, is a serious and growing threat to human health, posing challenges worldwide for the prevention and treatment of infections.^2,3^ A significant resistance mechanism in gram-negative bacteria is the acquisition and production of β-lactamases (Bla), enzymes that degrade β-lactam antibiotics.^4,5^ The rise of extended-spectrum-β-lactamases (ESBLs) that degrade all classes of β-lactams makes the development of new strategies to overcome Bla-mediated resistance an urgent medical need.

Available options to combat antibiotic-resistant infections include increasing the dosage of antibiotics, substituting for a narrow-spectrum agent targeted against the infection-causing pathogen, using an alternative last-resort antibiotic, and/or co-administering an antibiotic with a β-lactamase inhibitor. None of these approaches are satisfactory for all scenarios, can introduce further complications, and can even be counterproductive to antimicrobial stewardship efforts to minimize the spread of antibiotic resistance (**Fig. 1a**).^6^ Increasing the administered dose of β-lactam antibiotics has major acute and chronic impacts on the gut microbiome.^7,8^ Bacteria co-expressing multiple classes of β-lactamases make the use of Bla inhibitors challenging, especially since there are currently no clinically available inhibitors for metallo-β-lactamases (MBL).

**Fig. 1.**
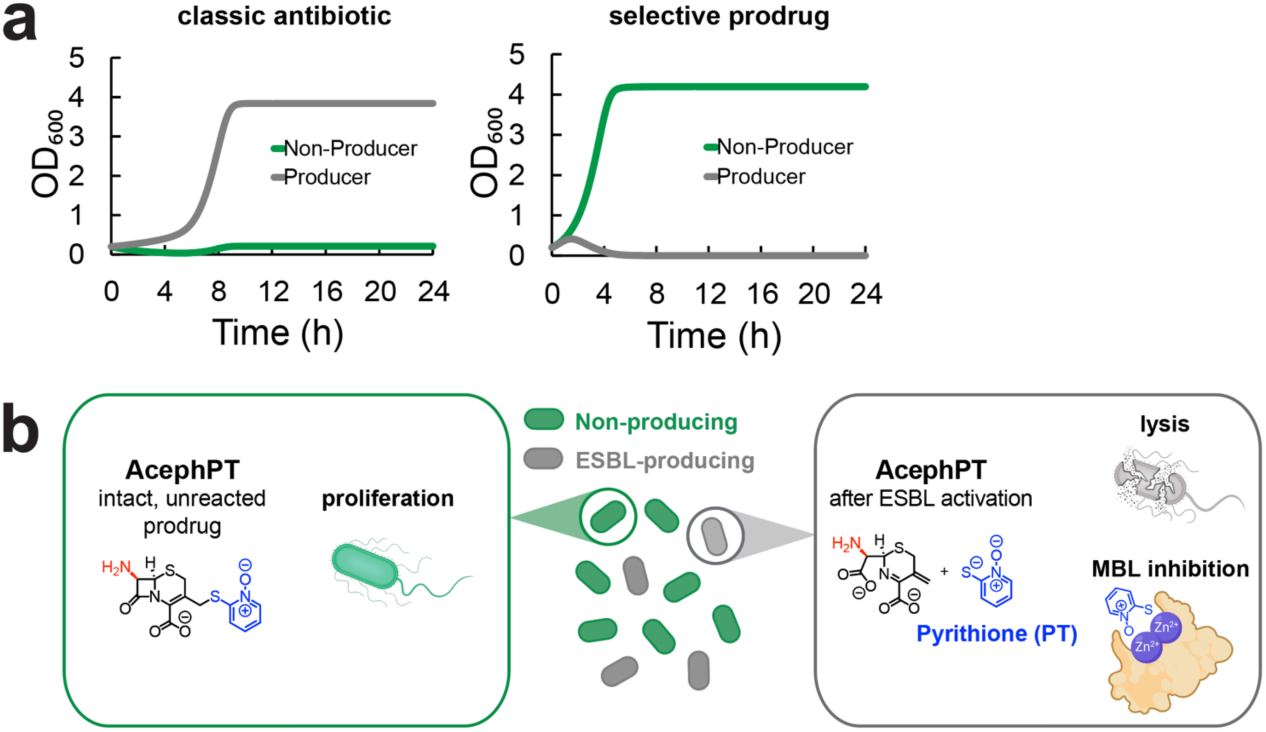
Antibacterial prodrugs to overcome the limitations of current antibiotics. **a**, Model of bacteria communal growth from different treatment strategies. Classic antibiotics (left) are ineffective at suppressing the growth of ESBL-producing bacteria, while there is collateral damage of ESBL-non-producing bacteria. A prodrug that is selectively activated by ESBL-producing bacteria (right) suppresses the growth of the resistant population while sparing the non-producing bacteria. **b**, Schematic of the prodrug AcephPT, which remains intact and ineffective at killing ESBL-non-producing bacteria (left) but is activated by ESBL-producing bacteria (right) leading to localized release of pyrithione, a bactericidal agent that also has metallo-β-lactamase inhibiting properties.

Given the limitations of current approaches, drug strategies that selectively suppress the proliferation of resistant populations among complex microbial communities are needed. Desirable agents would maintain the benefits of broad-spectrum coverage to increase the probability of clearing an infection of unknown composition, while minimizing collateral damage and the spread of resistance (**Fig. 1a**). Toward this goal, we introduce AcephPT, a prodrug intentionally designed to turn Bla reactivity into a vulnerability to achieve selective suppression of β-lactam resistant bacteria (**Fig. 1b**).

AcephPT is in the family of cephalosporins that release an active agent upon cleavage of the β-lactam ring upon hydrolysis by Bla enzymes.^9–16^ Like its previous generation prodrug PcephPT,^15^ AcephPT is designed to release pyrithione, a metal-binding antimicrobial linked to the cephalosporin core via one of its metal-binding atoms. As we previously showed for PcephPT, the conjugate has minimal metal-binding ability and cytotoxicity when pyrithione is linked in the prodrug state. However, in Bla-producing *Escherichia coli* this linkage is broken to release pyrithione, resulting in bacterial cell death. (**Fig. 1b**).^15^ Our mechanistic investigations revealed a second mode of action for PcephPT. Against strains expressing serine Blas, the in-situ released pyrithione was antimicrobial, while in carbapenem-resistant strains expressing the metallo-β-lactamase NDM-1, PcephPT also acted as an auto-inhibitor, with the cleaved pyrithione being retained at the Zn-active site of NDM-1, allowing NDM-1-producing bacteria to become sensitive to β-lactam antibiotics.^14,17^

While PcephPT was active against Bla-producing *E. coli*, it also retained conventional β-lactam activity and was cytotoxic against bacteria not producing Blas.^15^ This off-target activity is undesirable, as it would leave bacteria present in a healthy microbiome susceptible to suppression.

To achieve selective targeting of ESBL-expressing pathogens, we sought chemical modifications to the cephalosporin prodrug core that would favor activation by ESBLs while avoiding conventional β-lactam activity. Our strategy differs from traditional approaches that optimize chemical structures to prevent degradation by Blas. By purposefully designing a prodrug to invert the biased reactivity for Blas, we have forged AcephPT as a unique small molecule capable of selectively suppressing resistant, pathogenic bacteria even in the presence of bacteria that do not express β-lactamases. The design approach is guided by computational methods to identify molecular determinants capable of differentiating recognition by enzymes in ESBL producers and non-producers, enabling targeted activation and selective growth suppression of ESBL-producing *E. coli* in a mixed microbial community.

## Results

### Biasing prodrug recognition for ESBLs over PBPs

The enzyme targets of cephalosporins and other conventional β-lactam antibiotics are glycosyltransferase transpeptidases that crosslink peptidoglycans to mature the bacterial cell wall. Inhibiting these penicillin-binding proteins (PBPs) disrupts cell wall integrity, resulting in cell death by lysis. The activity of our first-generation prodrug PcephPT against Bla-non-producers suggests its structure makes it a reasonable substrate for PBPs. The suggested PBP recognition of PcephPT provides inadvertent off-target toxicity of non-resistant bacteria. Reactivity-based prodrugs that inhibit PBPs therefore lack the desired specificity for resistant, ESBL-producing bacteria.

We reasoned that achieving differential suppression of ESBL-producers vs non-producers would require differentiating the recognition of the small molecule for the active sites of ESBLs vs PBPs. Noting that clinically important cephalosporins contain functional groups appended to the β-lactam ring whereas Bla inhibitors like clavulanic acid and tazobactam do not, we hypothesized a bulky chemical structure, like the phenylacyl of PcephPT at the R^1^ site of the β-lactam ring (**Fig. 2a**), may facilitate interaction with PBPs but may not be necessary for engaging with Blas. To explore this structure-activity relationship, we used induced fit docking and molecular dynamics simulations to assess the engagement of select prodrugs with PBP3 and NDM-1. PBP3 was chosen because it is a known target of β-lactam antibiotics and is a component of the *E. coli* divisome essential for cell viability.^18,19^ NDM-1 is a metallo-β-lactamase with a di-zinc active site. It can hydrolyze nearly all β-lactam antibiotics likely due to a flexible binding pocket that accommodates a wide variety of β-lactam substrates.^20–22^ Because of the broad scope of β-lactam hydrolysis, NDM-1 and other MBLs are considered ESBLs for this study. For the prodrugs, we compared PcephPT with analogs containing smaller functional groups at the R^1^ site, specifically an acyl group for AcetamidocephPT and an amino group for AcephPT (**Fig. 2a**).

**Fig. 2.**
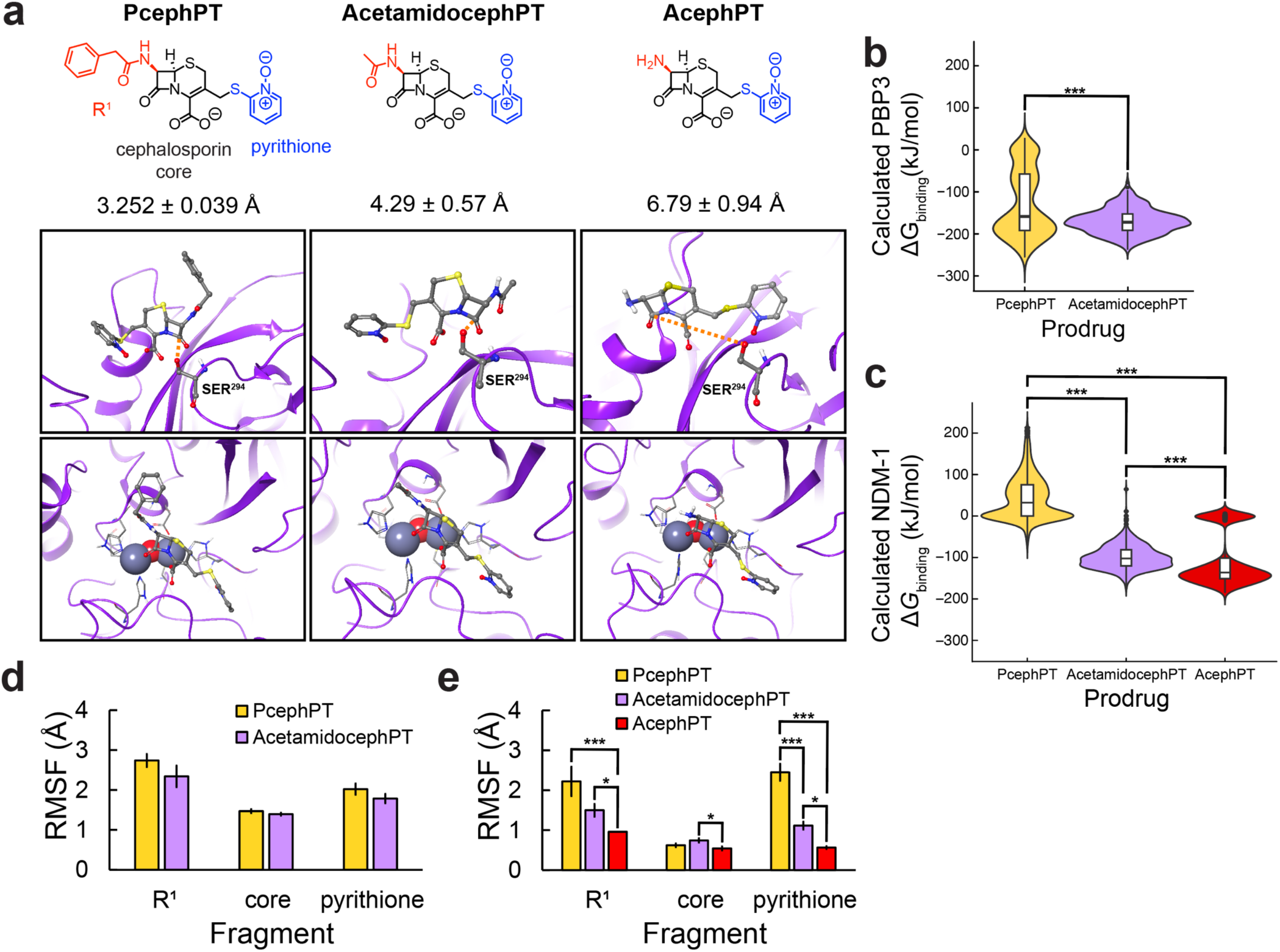
Computational modeling predicts AcephPT to be a worse substrate for PBPs and a better substrate for ESBLs. **a**, Representative induced-fit docking poses of PcephPT (left), AcetamidocephPT (middle), and AcephPT (right) to penicillin-binding protein 3 (PBP3, top panel), and the zinc metallo-β-lactamase NDM-1 (bottom panel). Distance between the catalytic serine residue of PBP3 (SER^294^) and the carbonyl carbon in the prodrug β-lactam ring is indicated by an orange dotted line, with values given for the mean ± SE from five poses in which the distance was shortest. Protein backbone depicted as purple ribbon, with SER^294^ and prodrugs depicted as ball-and-stick (showing polar hydrogens only; white = H, grey = C; blue = N; red = O, yellow = S), Zn-binding residues as tubes, and Zn and bridging hydroxide as spheres (grey = Zn). **b**, **c**, Violin plots overlaid with box-and-whisker plots showing the distribution of calculated Δ*G*_binding_ for prodrugs associated with PBP3. (**b**) and NDM-1 (**c**) across converged MD simulation (200–500 ns; *N* = 601 each group). Boxes are the interquartile ranges, whiskers are the upper and lower quartiles, center line is the median, points beyond the whiskers are included, potential outliers. *** p < 0.001 from univariate ANOVA with Tukey’s post-hoc test. **d**, **e**, RSMF values of heavy atoms in the given fragments of each of the prodrugs when associated and simulated with PBP3 (**d**) and NDM-1 (**e**). Bars are mean ± SE for the number of heavy atoms in each fragment. *** p < 0.001, * p < 0.05 from sequential univariate ANOVAs for data subdivision with Tukey’s post-hoc test or one-tailed t-test (R^1^ fragment, between groups).

Docking poses with PBP3 reveal notable differences in how AcephPT associates with the active site compared to PcephPT and AcetamidocephPT (**Fig. 2a**). Whereas the β-lactam cores of PcephPT and AcetamidocephPT are positioned in proximity to the catalytic SER^294^ (3.25 Å and 4.29 Å, respectively), the inverted orientation of AcephPT positions its carbonyl electrophile over 6.75 Å away from this serine, making it inaccessible for hydrolysis by PBP3. Hydrolysis of the β-lactam ring by PBP3 is a key step to producing a sterically trapped serine ester that covalently inhibits the protein to cause cell death.^23,24^ These induced-fit docking results indicate that PcephPT and AcetamidocephPT should be reasonable substrates for PBPs and predict that AcephPT would not be a physiologically relevant substrate for PBP3.

To gain insight into the interactions of the prodrugs with PBP3, molecular dynamic simulations were performed on PcephPT and AcetamidocephPT. AcephPT was not included in this analysis as it was deemed not to be a physiologically relevant substrate from the induced fit docking results. The free energies of binding were calculated from the ensemble of conformational snapshots from the converged molecular dynamic simulations over 200–500 ns (**Fig. S23-S27**). Comparison of the distribution of Δ*G*_binding_ of PcephPT and AcetamidocephPT to PBP3 indicates PcephPT’s sampled conformations lead to greater differences in association energy, with some conformations being more suitable for association with PBP3 (**Fig. 2b**). To further understand molecular features contributing to these differences in binding free energies, atom RMSF values for each prodrug were computed to quantify how far individual heavy atoms of the prodrug move away from the enzyme backbone across the simulation time. Increases in atom RMSF values translate to increased atom movement away from the protein across sampled conformations, a determinant of relative stability – the more movement, the lower the stability. Following a technique from fragment-based drug design,^25,26^ the contribution of individual fragments (the R^1^ functionality, the cephalosporin core, or the pyrithione leaving group) to the movement of the whole molecule within the active site were averaged and compared (**Fig. 2d**). No statistical significance between the fragment contributions of PcephPT and AcetamidocephPT was found, indicating both prodrugs have similar stability within the active site of PBP3.

In contrast with results from PBP3, induced fit docking poses of the test molecules with NDM-1 did not reveal any notable recognition differences across PcephPT, AcetamidocephPT, or AcephPT, suggesting all three compounds would be suitably recognized as substrates for this enzyme (**Fig. 2a**, bottom). This finding is consistent with reports of the broad substrate recognition of NDM-1.^20,21,27^ Molecular dynamic simulations performed with PcephPT, AcetamidocephPT, and AcephPT associated with NDM-1 yielded Δ*G*_binding_ values showing AcephPT to be a more favorable substrate for NDM-1 compared to PcephPT and AcetamidocephPT (**Fig. 2c**). The RMSFs of the prodrugs’ fragments with NDM-1 reveal the phenylacyl group of PcephPT experiences the largest fluctuation of the R^1^ fragments with NDM-1 (**Fig. 2e**). Notably, replacing the phenylacyl R^1^ with an acyl group in AcetamidocephPT or an amine in AcephPT suppresses the fluctuations at these positions. Additionally, the total stability of AcetamidocephPT and AcephPT improves compared to PcephPT, as a change in the R^1^ fragment corresponds with stabilization of their respective pyrithione fragments (**Fig. 2e**). Taken together, these computational results provide a rationale for modifying the R^1^ position of cephalosporin prodrugs with smaller groups to retain desirable recognition by ESBLs while suppressing recognition by PBPs. AcephPT and AcetamidocephPT were therefore synthesized to experimentally test these predictions. Synthetic details and characterization data are available in the Supplementary Information.

### AcephPT selectively suppresses ESBL-producing bacteria in clonal populations

We tested PcephPT, AcetamidocephPT, and AcephPT against a Bla-non-producing strain of *E. coli* MG1655 and MG1655 expressing different Bla enzymes. The panel of Blas consisted of serine-Blas (SBLs): OXA-1, TEM-1, CMY-2 and CTX-M-1, as well as zinc-containing MBLs: NDM-1, VIM-2, and IMP-1. Among this series, CTX-M-1, NDM-1, VIM-2, and IMP-1 (**Fig. 3a-c**, greens) are classified as ESBLs and carbapenemases, presenting promiscuous β-lactam recognition allowing bacteria expressing these enzymes to confer resistance to a wide range of antibacterials, including last-resort carbapenems.^20,22,28–32^ OXA-1, TEM-1, CMY-2 are also clinically significant Blas, but their scope of β-lactam recognition is narrower than the ESBLs and carbapenemases. (**Fig. 3a-c**, greys).^33–35^

**Fig. 3.**
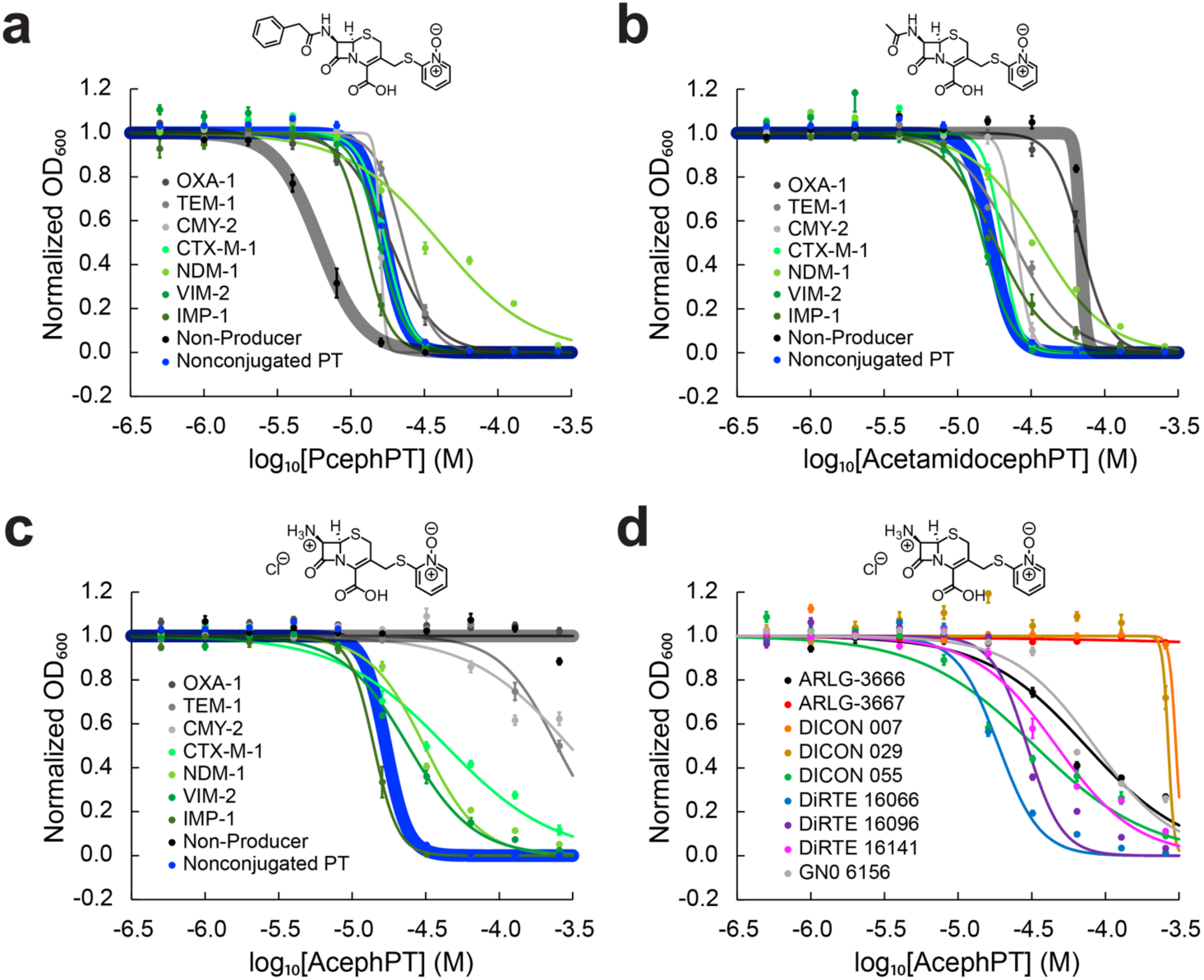
AcephPT selectively suppresses ESBL-producing lab variety bacteria and clinical isolates. Dose-response curves of **a**, PcephPT **b**, AcetamidocephPT, and **c**, AcephPT treatment of *E. coli* K-12 MG1655 engineered to express the indicated Blas, with a non-producer shown in translucent black and those classified as ESBLs in green. Treatment of pyrithione as a non-conjugated small molecule shown in blue. **d**, Dose-response curves of AcephPT against a panel of clinical isolates producing Blas (see **Table S1** for species and Blas produced). Points and error bars are mean ± SE from 3 biological replicates of 4 technical replicates (*N* = 12 each group). For Nonconjugated PT, points and errors bars are mean ± SE from 3 biological replicates from each bacterial strain tested (*N* = 24 each group). Fitted lines are logistic functions determined from non-linear regression.

As a positive control, all our engineered bacteria were challenged by free pyrithione, for which EC_50_ values ranged from 15–20 µM. (**Fig. S28**) The narrow range in pyrithione EC_50_ against each of these strains indicates Bla expression does not impact pyrithione’s activity. The average of these bacterial growth dose-response curves is depicted as the blue curve for Nonconjugated PT in **Fig. 3a-c** and represents the toxicity threshold for the active agent. In Bla-producing strains, pyrithione-conjugated prodrugs achieving EC_50_ values near this pyrithione toxicity threshold of 17 µM indicates good drug activation by Blas.

PcephPT inhibited growth of all strains tested, with EC_50_ values ranging from 6–40 µM (**Fig. 3a**). The fact that the Bla-non-producing strain was the most sensitive while the NDM-1-producing strain was the least sensitive highlights PcephPT’s lack of desired selectivity for Bla-producing bacteria.

Removing the phenyl moiety from the R^1^ functionalization to produce AcetamidocephPT resulted in ten-fold decreased activity against Bla-non-producing bacteria compared to PcephPT (**Fig. 3b**). Although AcetamidocephPT was least toxic to the strain producing the penicillinase OXA-1, for all other Bla-producing strains tested AcetamidocephPT retained a similar level of activity to PcephPT, with EC_50_ values ranging from 15–36 µM.

Following the trend of decreasing chemical space at the R^1^ position of the prodrug core, removal of the acyl group on AcetamidocephPT leaves the amine functionality of AcephPT. When tested against our Bla-non-producing bacteria, AcephPT was remarkably non-toxic (**Fig. 3c**), as no growth suppression was observed for the concentrations tested. The strains producing the Blas OXA-1, TEM-1, and CMY-2 were also less sensitive to AcephPT, with EC_50_ thresholds not being reached by the concentrations tested (EC_50_ > 256 µM). The strains producing the ESBLs CTX-M-1, NDM-1, VIM-2 and IMP-1, however, were sensitive to AcephPT, with EC_50_ values ranging from 14–43 µM. The non-ESBL Bla-producing variants do not recognize AcephPT, while the more promiscuous ESBL-producing bacteria recognize AcephPT and remain sensitive, corroborating the predictions from our molecular modeling studies with PBP3 and NDM-1. Notably, AcephPT is active against Bla types that produce the greatest amount of resistance to current antibiotics.

Given AcephPT’s differential activity against our panel of engineered Bla-non-producing and ESBL-producing bacteria, we further tested it across a panel of clinical isolates acquired from patients treated in the United States. Sites of infection for the clinical isolates include joint and blood infections. The panel was selected to represent different Bla types and bacterial species (**Fig. 3d**). Details on species and Blas expressed in each clinical isolate is available in **Table S1**. For all strains except the ARLG 3667 *Pseuodomonas aeruginosa*, the positive control of free pyrithione was effective at suppressing the growth of the Bla-producing clinical isolates, with EC_50_ ranging from 11–37 µM (**Fig. S29**). The Bla-producing *P. aeruginosa* strain was not sensitive to free pyrithione at the concentrations tested (EC_50_ > 256 µM).

AcephPT’s EC_50_ values determined against *E. coli*, *Klebsiella pneumoniae*, and *Enterobacter cloaca*e clinical isolates (**Fig. 3d**) were similar to those found for our Bla-producing engineered *E. coli* (**Fig. 3d**). The exceptions to this trend were ARLG 3667, the *P. aeruginosa* isolate that was also not susceptible to free pyrithione, and DICON isolates 007 and 029. The observation that AcephPT was ineffective against DICON 007, an *E. coli* isolate that produces TEM-1, is consistent with the poor activity found against our engineered strain producing TEM-1, which is not considered an ESBL (**Fig. 3c**). DICON 029 has been annotated to coproduce CTX-M-15, an ESBL, and OXA-48, not an ESBL. The weak activity of AcephPT against this strain may suggest additional mechanisms of resistance. Overall, AcephPT suppressed the growth of clinically relevant bacteria that are otherwise resistant to broad-spectrum antibiotics.

### AcephPT is hydrolytically activated exclusively by ESBL-producing *E. coli*

Results from the dose-response curves are consistent with a mechanism in which the β-lactam structure of AcephPT is stable to non-specific hydrolysis, yet susceptible to ESBL-triggered release of pyrithione as the active cell-killing agent. To verify the molecular details of drug activation in cells, we adapted^36^ and designed a time-course ^1^H NMR spectroscopy experiment to profile the activation of AcephPT in our engineered ESBL-non-producing and ESBL-producing *E. coli* MG1655. Proton resonances of the prodrug are observed and monitored for consumption by the expressed ESBL. Changes in resonance frequencies indicate a new species is being produced, such as the release of pyrithione from the conjugate. As shown in **Fig. 4a**, AcephPT remained intact with no hydrolysis or other decomposition being observed upon incubation with the Bla-non-producing strain across the incubation time. In contrast, notable changes in the resonance peaks of AcephPT are observed upon incubation with the strain that expresses the SBL CTX-M-1 (**Fig. 4b**). As the starting prodrug’s resonances decrease in intensity, resonances corresponding to free pyrithione increase in the aromatic region (8.4–6.9 ppm) with the identical intensity change. Additionally, the appearance of alkenyl resonances in the 5.9–5.6 ppm region is indicative of alkene formation within the hydrolyzed prodrug fragment (see **Fig. 1b**), which is mechanistically anticipated to form during the release of pyrithione.^10,11,14^ Similar resonance changes were observed when AcephPT was incubated with our engineered IMP-1-producing (**Fig. 4c**) and NDM-1-producing strains (**Fig. S30**).

**Fig. 4.**
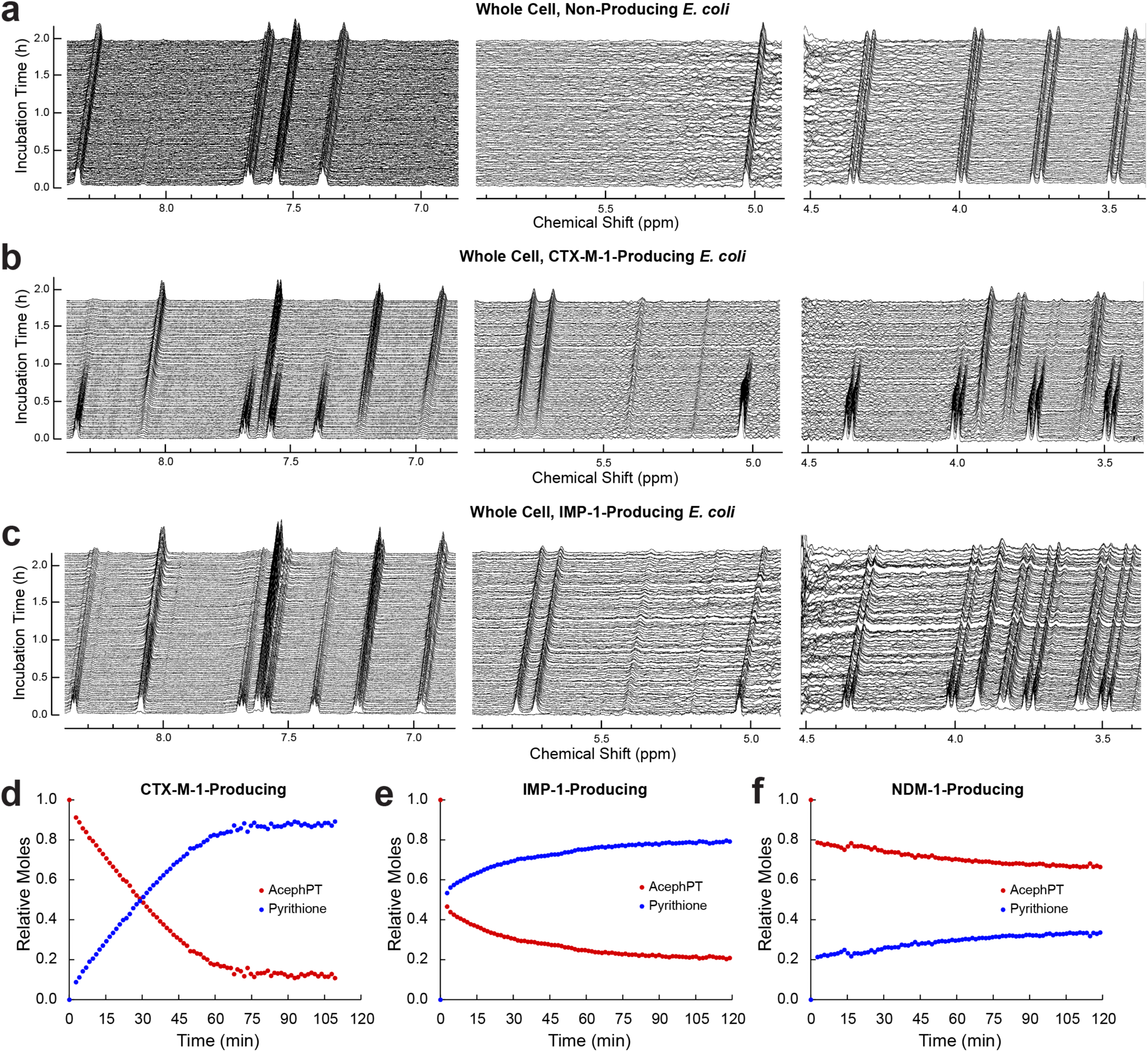
AcephPT is selectively activated by ESBL-producing *E. coli*. Time-course NMR spectra profile the chemical changes of the prodrug AcephPT incurred upon incubation with whole cell *E. coli* at OD_600_ = 2.4–2.6 for **a**, Bla-non-producing **b**, CTX-M-1-producing, and **c**, IMP-1-producing *E. coli*. The unchanged resonances in **a** show stability of AcephPT in the non-producer strain. **d**, **e**, **f**, Speciation curves for the enzymatic transformation of AcephPT to pyrithione for CTX-M-1, IMP-1, and NDM-1-producing *E. coli*, respectively, from integrations of the individual NMR spectra. CTX-M-1-producing *E. coli* hydrolyzes all available AcephPT. IMP-1- and NDM-1-producing *E. coli* hydrolyze an initial amount of AcephPT rapidly before the reaction stalls. Each point is an average of the integrations for resonances in the aromatic region (6.9–8.5 ppm) for the corresponding species scaled to an external standard of AcephPT in the absence of *E. coli* (*t*_0_).

When AcephPT was incubated with our CTX-M-1-producing strain, the intensity of the resonances observed for the prodrug’s consumption matched the intensity of the peaks corresponding to free pyrithione (**Fig. 4d**), indicating stoichiometric conversion from prodrug to active drug. Additionally, conversion of the prodrug to active drug went to completion in this strain. When incubated with our IMP-1-producing strain, AcephPT hydrolysis was initially more rapid than with the CTX-M-1-producing strain, as more than 0.5 molar equivalents of AcephPT were converted to free pyrithione by the acquisition of the first spectrum (**Fig. 4e**). This activity plateaus, as the hydrolysis of AcephPT by the IMP-1-producing strain significantly slows down, leaving 0.2 molar equivalents of the prodrug still intact after the two-hour incubation. The stalling of prodrug activation was also captured in our NDM-1-producing strain, which activated 0.2 molar equivalents of AcephPT before acquisition of the first spectrum, and only hydrolyzed another 0.1 molar equivalents across the two-hour incubation (**Fig. 4f**). This stalling of prodrug activation is reminiscent of our previous work with PcephPT, in which PcephPT treatment led to inhibition of NDM-1.^14^

Holistically, these in-cell ^1^H NMR experiments demonstrate the excellent stability of AcephPT in aqueous buffered and cellular environments at physiological temperature in the absence of Bla-producing bacteria. Furthermore, they validate the release of pyrithione upon hydrolysis of AcephPT (**Fig. 4d-f**), and that prodrug hydrolysis and release of active pyrithione is specific for ESBL-positive bacteria.

### AcephPT selectively suppresses a drug-resistant clinical isolate in a pairwise mixed microbial environment

To determine the selectivity of the prodrug in a communal setting, we created mixed cultures of 1:999, 1:99, and 1:9 proportions of ESBL-producing to non-producing strains (**Fig. 5a**). The ESBL-producing strain was the pathogenic DiRTE 16096 *E. coli* clinical isolate that expresses NDM-1 (AcephPT EC_50_ = 29 µM). To model a Bla-non-producing “commensal” strain, we used an engineered laboratory Top10 *E. coli* that expresses mCherry and lacks β-galactosidase (AcephPT EC_50_ > 256 µM, see **Figure S31**). The mixed cultures were prepared from individual overnight cultures equalized to an OD_600_ of 1.0. The cultures were diluted 100-fold and were untreated or treated with cefotaxime (a third-generation β-lactam) or AcephPT in 96-well plates. While the mixtures were incubating over 24 hours, OD_600_ readings were taken to indicate total culture density of the mixture (**Fig. 5b**) and fluorescence intensity measurements (Ex_586_/Em_620_) were taken to monitor mCherry expression (**Fig. 5c**), a distinct reporter of growth of the Top10 Bla-non-producing strain.

**Fig. 5.**
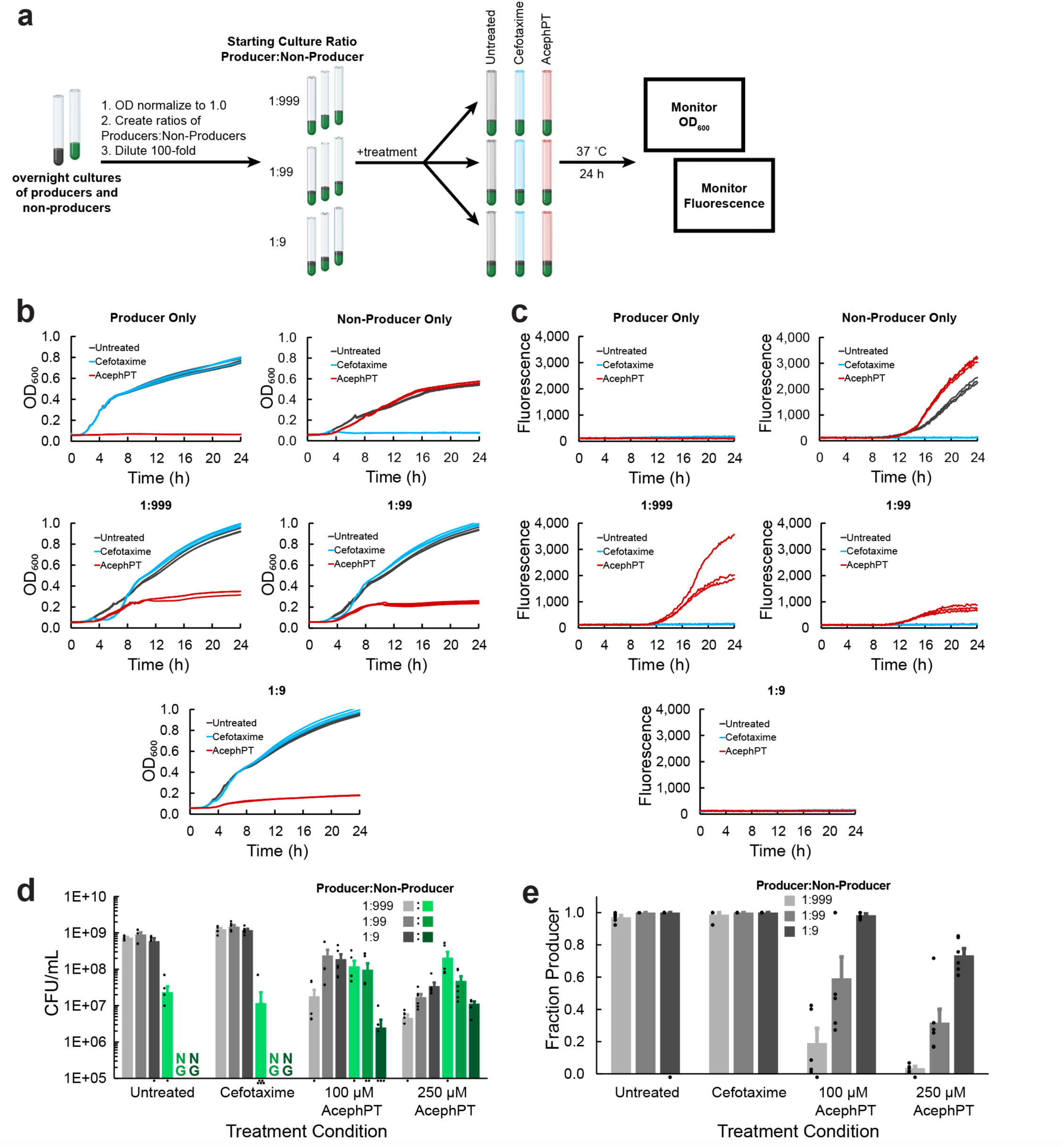
In communal mixtures, AcephPT suppresses the growth of ESBL-producing, resistant *E. coli* while recovering the growth of non-producing, non-resistant *E. coli*. **a,** Schematic of communal experimental design. Different producer:non-producer ratios were prepared from individual overnight starter cultures before being treated. Conditions were monitored optically for **b**, culture growth by OD_600_ and **c**, for expression of mCherry by fluorescence (Ex_586_/Em_620_) for 24 h. **b**, Growth curves of cultures for each producer:non-producer ratio under each treatment condition. The producer strain (top left) is sensitive to AcephPT treatment but not cefotaxime. The non-producer strain (top right) is sensitive to cefotaxime treatment but not AcephPT treatment. The producer strain grows faster than the non-producer strain. All communal mixtures show no change in growth upon cefotaxime treatment. Treatment with AcephPT shows growth suppression for each mixture, with the greater starting ratios of producers being more sensitive to AcephPT. Each curve is a biological replicate, showing 3 for each group. **c**, Fluorescence measurements taken in tandem with culture growth curves to monitor expression of mCherry. No fluorescence is detected for producers (top left), as they do not express mCherry. For the non-producers (top right), fluorescence increases over time in the untreated and AcephPT conditions, but not the cefotaxime-treated conditions, as the non-producers are sensitive to this antibiotic (**b**, top left). All communal mixtures show no change in fluorescence in the untreated and cefotaxime-treated conditions, indicating culture take over by the producers. Fluorescence for the AcephPT treated conditions are present in the 1:999 and 1:99 mixtures, indicating growth recovery of the non-producer. Each curve is a biological replicate, showing 3 for each group. **d**, CFU/mL enumerations of viable producers (greys) and non-producers (greens) for different treatment conditions determined from blue/white (producers/non-producers) screening. Cefotaxime treatment results in no change culture ratio compared to untreated, indicating it cannot combat the ESBL-producing population. AcephPT suppresses growth of the ESBL-producing population at all starting culture ratios and recovers growth of the non-producing population. AcephPT’s activity is dose-dependent in communal environments, as changes in culture composition are more extreme from 100 to 250 µM. Bars are average ± SE from 3 biological replicates of 2 technical replicates (*N* = 6 each group). Black circles are individual data points; points under the axis indicate no viable growth for that replicate. Data areas with NG labels had no viable cell growth for that condition. **e**, Fraction of ESBL-producing *E. coli* determined from viable cell enumerations (**d**) by dividing raw counts of producers by the total for each ratio/treatment condition. AcephPT suppresses the growth of ESBL-producing *E. coli*. Bars are average ± SE from 3 biological replicates of 2 technical replicates (*N* = 6 each group). Black circles are individual data points; points under the axis have an undefined value as no viable producers or non-producers were found for the given replicate.

The ESBL-producing strain is sensitive to AcephPT treatment, but not cefotaxime (**Fig. 5b**, top left). In contrast, the Bla-non-producing strain (**Fig. 5b**, top right) is sensitive to cefotaxime but not AcephPT. The ESBL-producer grows faster than the non-producer, as the culture density is greater in the untreated condition for the ESBL-producer. For the communal mixtures (**Fig. 5b**, middle and bottom) treated with cefotaxime, there is no change in culture density from their respective untreated conditions. Treatment with AcephPT, however, results in suppressed growth for each mixture compared to untreated or cefotaxime-treated conditions (**Fig. 5b**, middle and bottom). Decreases in overall culture density from AcephPT treatment correlates with the starting proportion of producers – as the starting culture ratio of producers increases, the AcephPT treated communal culture grows less.

The fluorescence measured in tandem with culture growth curves were used to monitor expression of mCherry and therefore growth of the non-producer. No fluorescence was detected for producers (**Fig. 5c**, top left), as they do not express mCherry. For the non-producers (**Fig. 5c**, top right), fluorescence increased over time in the untreated and AcephPT conditions, correlating well with increases in the cell density curves (**Fig. 5b**, top right). The cefotaxime-treated non-producer condition presented no increase in fluorescence, validating that non-producers are sensitive to this antibiotic (**Fig. 5b**, top right). For the non-producers, the lack of florescence from cefotaxime treatment but increase in florescence from AcephPT treatment means the non-producers are sensitive to cefotaxime but not sensitive to AcephPT. None of the communal mixtures showed any increase in fluorescence in the untreated and cefotaxime-treated conditions, indicating culture takeover by the producers. This producer takeover highlights the importance of bacterial growth dynamics in complex environments with and without exposure to classic antibiotics. In contrast, fluorescence for the AcephPT treated conditions increase in the 1:999 and 1:99 mixtures, indicating growth recovery of the non-producer (**Fig. 5c**, middle). Together, these data indicate AcephPT is targeting the resistant bacteria and allowing for recovery of the ESBL-non-producing bacteria.

While the fluorescence readout of mCherry expression allows for the rapid screening of ESBL-producing and ESBL-non-producing pairs, it does not quantify the proportion of viable bacteria from each strain. To enumerate colony-forming units (CFUs) of viable bacteria, treatment and mixture combinations were grown for 24 hours, diluted, and spotted onto agar plates containing X-Gal and IPTG (**Fig. 5d**). Since the *E. coli* Top10 engineered non-producer strain does not have β-galactosidase, these colonies appear white, allowing for differentiation from the blue colonies of the clinical isolate producer strain during blue-white screening to quantify viable cells of each type.^51^ In the 1:99 and 1:9 mixtures, no growth of non-producer bacteria was observed in the conditions treated with vehicle or cefotaxime. This result recapitulates those depicted in **Fig. 5b-c**; the growth rate of the producers allows for its takeover of the communal environment. Occurring in starting ratios as low as 1:99, this takeover again highlights the importance of bacterial growth dynamics in mixed cultures. In contrast, AcephPT significantly suppressed the producer population in all mixtures, while encouraging the growth of the non-producers (**Fig. 5d**). AcephPT selectively suppresses the ESBL-producing strain in a manner depending on both the starting culture ratio and the dose. The proportion of pathogenic cells in the mixed population treated with AcephPT was reduced up to 100-fold compared to untreated and cefotaxime-treated mixtures.

To highlight the suppression of the producer, the fraction of producers remaining after treatment was determined for each treatment and mixture combination (**Fig. 5e**). Cefotaxime presented no suppression of the producer population, being no different from the untreated conditions. AcephPT, in contrast, suppressed the growth of ESBL-producing *E. coli*. These key data show prodrug AcephPT preferentially suppresses the producer population in each mixed microbial environment, allowing reversal of the fraction of pathogenic vs commensal bacteria, in favor the ESBL-non-producing bacteria.

These results reinforce that treatment with a classic antibiotic such as cefotaxime is not only ineffective for combating a drug-resistant mixture but selects for growth of the drug-resistant bacteria. AcephPT overcomes this limitation by exploiting the resistance mechanism to suppress the growth of the ESBL-producer and promote the growth of the non-producer.

## Discussion

Antibiotic use has counter-productively created selection conditions for spreading extended spectrum β-lactamases (ESBLs), which have promiscuous recognition and enhanced kinetic activity to hydrolyze and inactivate β-lactam antibiotics. Prodrug AcephPT was designed to take advantage of these properties by being purposefully hydrolyzed, and therefore activated, by ESBLs to become cytotoxic. Remarkably, AcephPT does not affect Bla-non-producing bacteria because penicillin-binding protein (PBP) recognition was diminished by its molecular design. AcephPT demonstrates a first-in-class ability to suppress ESBL-producing, pathogenic bacteria while simultaneously allowing ESBL-non-producing bacteria to recover. By turning ESBL expression into a weakness, reactivity-based prodrugs like AcephPT may improve antimicrobial stewardship by selectively suppressing resistant populations.

Key to the design strategy that led to AcephPT was de-engineering the susceptibility of non-ESBL-producing strains to what is otherwise a classic cephalosporin antibiotic scaffold. The unabated growth of non-ESBL-producing *E. coli* in the presence of increasing concentrations of AcephPT substantiated this idea, which was validated by whole cell NMR experiments confirming the molecular stability of AcephPT against unintended hydrolysis reactions or degradation in a cellular context of non-ESBL-producing *E. coli*. Furthermore, the NMR experiments were critical for characterizing the chemical products of prodrug activation by ESBL producers. This characterization is important for establishing the mechanism of action, as hydrolysis of other reactivity-based β-lactam compounds has been shown to occur without release of the leaving group.^10^

Quantifying the release of pyrithione by bacteria expressing different ESBLs further revealed that rates of release and level of autoinhibition vary based on the expressed ESBL. To be amenable to the sensitivity of the NMR experiment, the cell densities and concentrations of compound used for these experiments were significantly greater than those used in microculture cell assays. Under these conditions, the in-situ NMR experiments revealed auto-inhibition exclusively by strains expressing MBLs. Hydrolysis of AcephPT by the SBL CTX-M-1 was steady until the prodrug was completely consumed. When incubated with the MBL strains expressing IMP-1 or NDM-1, there was an initial burst of pyrithione released before hydrolysis significantly slowed down. Our previous work with first-generation prodrug PcephPT revealed mixed activity of pyrithione as both an antimicrobial and an inhibitor of the MBL NDM-1, which was attributed to pyrithione’s metal binding activity.^14^ AcephPT appears to maintain this paired activity, as the stalling of enzymatic activity is likely due to sufficient pyrithione being released to inhibit both of the MBLs tested here. Comparatively, IMP-1 was inhibited less than NDM-1, as IMP-1 turned over more AcephPT before activity stalled.

These observations raise an interesting conundrum, as autoinhibition by reactivity-based prodrugs could undercut the mechanistic advantage providing their selectivity. When the active agent can serve as both cytotoxic agent and inhibitor, as it does for AcephPT, the interplay of the rate of release, the EC_50_ of the cytotoxic agent, and the enzyme inhibition constant will influence the overall capability of generating sufficient biocide without prematurely inhibiting the enzyme that releases it. A further consideration is the possibility of collateral damage by the released agent, which if overproduced could become available to inflict nonspecific damage to cells within its diffusion reach. Under these scenarios, the dynamics of autoinhibition of MBLs may be favorable for self-tempering such an outcome.

The observation that MBL-expressing strains were the most sensitive of the strains tested to growth inhibition by AcephPT (**Fig. 3c**) suggests the conditions in the microdilution experiments were such that sufficient pyrithione was released to reach efficacious levels to inhibit growth of the lab strain monocultures without reaching auto-inhibition levels. In the mixed microbial cultures, the additional factor of population dynamics comes into play.

As our relatively simple two-strain pairwise mixtures show, even a very small percentage of a faster-growing pathogen readily outcompetes to take over the whole culture (**Fig. 5b**). While pyrithione was found not to be bactericidal to the clinical isolate producer strain used in the mixed microbial experiment, it was bactericidal to the non-producer strain at 250 µM (**Fig. S31**). This difference in cidality between the two strains explains why the producer was not fully eliminated in our mixed microbial model and implies pyrithione was not over-produced to reach collateral bactericidal levels available to the non-producer. Although the producing strain was not eliminated, the reduced growth of the strain is noteworthy, and the recovery of the slower-growing non-producer, most evident at the lowest ratios of producer, is remarkable.

Our work illustrates AcephPT as a potential targeted therapeutic agent against drug-resistant bacteria, opening the door for further development of antibacterials that are selectively recognized by bacteria producing ESBLs. We demonstrate the pharmacophore pyrithione can selectively suppress ESBL-positive pathogenic bacteria by its conjugation to a cephalosporin, with modification to its β-lactam core that makes it a worse PBP substrate while maintaining activity with ESBLs. The enzymatic activity generating drug resistance (i.e. cleavage of β-lactam drugs by Blas) can be hijacked to release antimicrobial compounds that selectively suppress the pathogens harboring these resistance enzymes, while sparing others. With this design strategy we can push back against evolutionary trends to curb β-lactamase derived resistance while using it to our advantage to selectively release active pharmacophores to pathogenic bacteria.

## Materials and Methods

### General Chemical Information

Synthetic procedures and characterization details for all compounds can be found in the Supplementary Information. All reagents and solvents were purchased from commercial sources (Millipore Sigma, Oakwood Chemical, Fischer Scientific, AA Blocks, Combi Blocks) and used as received. Silica gel (230–400 mesh) was used as a stationary phase for column chromatography. Eluent details are listed with the associated compound in the Supplementary Information. TLC was performed on silica gel 60 F_254_.

### NMR Spectroscopy

^1^H, ^13^C{^1^H}, DEPT-90, and DEPT-135 NMR data were acquired on a Bruker Ascend 500 MHz instrument at ambient temperature. Chemical shift values are reported in ppm with coupling constants in Hz. Abbreviations for multiplicity are: br = broad singlet, s = singlet, d = doublet, t = triplet, q = quartet, dd = doublet of doublets, dt = doublet of triplets, ABq = H_A_H_B_ quartet, m = multiplet. The solvent signal for each spectrum was used for axis calibration. NMR data were processed using a Bruker TopSpin academic license. Resonance assignments and spectra for ^1^H and ^13^C{^1^H} spectra are available in the Supplementary Information.

### Mass Spectrometry

Liquid chromatography/mass spectrometry (LC/MS) analyses were acquired on an Agilent 1260/6460 instrument. Positive-ion mass spectra were acquired in full-scan mode over the range of 100–2500 m/z using the following source parameters: gas temperature 300 °C, gas flow 5 L/min, nebulizer pressure 45 psig, VCap 3500 V, and fragmentor voltage 135 V. HIC HPLC separations were achieved on a Phenomenex Luna C18(2) column (100 mm x 2 mm, 3 µm, 100 Å) using a linear gradient of mobile phase B in A, a flow rate of 0.5 mL/min, and a column temperature of 40 °C. Mobile phase A was prepared by combining 400 mL ultrapure water with 12 mL methanol and 1.2 mL formic acid. Mobile phase B was prepared by combining 400 mL acetonitrile with 12 mL ultrapure water and 1.2 mL formic acid. The gradient program included an initial hold at 0 % B for 2 min, linear increase to 60 % B from 2–9 min, linear increase to 100 % B from 9-10 min, hold at 100 % B from 10-12 min, linear decrease to 0 % B from 12–13 min, final hold at 0 % B from 13-15 min for a total run time of 15 min. HILIC HPLC separations were achieved on a SeQuant ZIC-HILIC column (100 mm x 2.1 mm, 3.5 µm, 100 Å) using a linear gradient of mobile phase B in A, a flow rate of 0.5 mL/min, and a column temperature of 40 °C. Mobile phase A was 5 mM ammonium formate at pH 2.5, prepared by dissolving ammonium formate in ultrapure water and adjusting pH with formic acid. Mobile phase B was prepared by combining 400 mL acetonitrile with 12 mL ultrapure water and 1.2 mL formic acid. The gradient program included an initial hold at 95 % B for 2 min, linear decrease to 40 % B from 2–9 min, linear decrease to 5 % B from 9-10 min, hold at 5 % B from 10-12 min, linear increase to 95 % B from 12–13 min, final hold at 95 % B from 13–15 min for a total run time of 15 min. In addition to MS detection, the DAD was used to acquire a UV chromatogram at 220 nm, 254 nm, and 280 nm. Samples were analyzed using a 2 µL injection volume. Processed raw data files were exported from Agilent MassHunter and plotted in Microsoft Excel. UV chromatograms and mass spectra of isolated compounds are available in the Supplementary Information.

High-resolution mass spectra (HRMS) were acquired on an Agilent 1200/6224 instrument. The mass spectrometer was equipped with a Dual ESI source, and accurate mass data were obtained by internal calibration (reference ions 121.050873 and 922.009798 m/z) using a secondary nebulizer to continuously deliver the reference solution. Positive-ion mass spectral data were acquired in full-scan mode over the range of 100–3200 m/z using the following source parameters: gas temperature 325 °C, gas flow 11 L/min, nebulizer pressure 33 psig, VCap 3500 V, and fragmentor voltage 220 V. Samples were analyzed using a 1-5 µL injection volume. HRMS results of isolated compounds are available in the Supplementary Information.

### Communal Growth Model

For simulations shown in Figure 1, ordinary differential equations were formulated to model the qualitative dynamics of a bacterial system with ESBL-producing and Bla-non-producing subpopulations responding to an antibiotic or a prodrug. The dynamics were formulated as the interactions between six main components: ESBL-producing population density (*n_r_*), Bla-non-producing population density (*n_s_*), nutrient level (*s*), antibiotic concentration (*a*), extracellular ESBL concentration (*b*), prodrug concentration (*p*), and released drug concentration (*e*). In this model, ESBL production extracts a fitness cost (*α*) and grants resistance to the antibiotic (*β*) relative to the Bla-non-producing population. Nutrient is consumed by growth and released incompletely (*ξ*) with lysis. Antibiotic degradation and prodrug cleavage are mediated either by extracellular ESBLs (κ_b_, κ_p_) which are released on lysis, or living ESBL producers (φ, σ). All simulations were conducted in MATLAB R2023b. Details of the differential equations and the model can be found in Supplementary Information.

### Molecular Modeling

Using Spartan 20 (https://wavefun.com/), an equilibrium geometry calculation was performed on each β-lactam small molecule. The method was as follows: Density-function theory (DFT) = B3LYP 6-311+G**, solvent = conductor-like polarizable continuum model (CPCM), dielectric = 78.3 (water). The DFT-optimized β-lactam structures were exported to SDF format and transferred to Maestro (Schrödinger Inc.). The Ligand Preparation task was launched. No filter criteria were used, OPLS4 force field was used, target ionization set to 7.4 ± 3.0 using Epik and metal binding states were included with the original state of the ligand also requested to be kept. Additionally, ligands were desalted and tautomers were generated. Specified chirality of the small molecule was retained, but a max of 32 stereoisomers were allowed to be prepared if possible.

X-ray diffraction crystal structure of PBP3 complexed with hydrolyzed imipenem [Protein Data Bank (PDB) 3PBQ]^37^ was imported to Maestro. All waters were removed from the structure entry. X-ray diffraction crystal structure of NDM-1 complexed with hydrolyzed ampicillin [Protein Data Bank (PDB) 5ZGR]^38^ was imported to Maestro. The structure is a non-covalent dimer of NDM-1; protein, ligands, waters, and metal ions associated with structure B were removed from the structure entry. Additionally, glycerol and all waters except the active site hydroxide (HOH 412) were removed from the structure entry. The Protein Preparation Workflow task was launched. Preprocessing was performed with the following parameters: fill in missing side chains; assign bond orders, using Combined Chemical Dictionary (CCD) database; replace hydrogens; create zero-order bonds to metals; create disulfide bonds; fill in missing loops using Prime; sample water orientations, use crystal symmetry, minimize hydrogens of altered species, use PROPKA with pH 7.4; restrained minimization was then performed, converging heavy atoms to root mean square deviation of 0.3 Å using OPLS4; waters were removed 3.0 Å beyond het groups. All default positions for alternate residue positions were committed. For the prepared PBP3, the ester bond between the catalytic serine residue (SER^294^) and hydrolyzed imipenem was broken. For the prepared NDM-1, the additional hydrogen atom on the active site water was deleted to form the appropriate active site hydroxide. Additionally, the N-H bond formed on ampicillin was deleted.

With the proteins and β-lactam small molecules prepared, Induced Fit Docking was launched to dock the small molecules to PBP3 or NDM-1. The receptor box was centered about the complexed ligand from the parent X-ray diffraction crystal structure. Prime refinement was performed to optimize residues within 12.0 Å of the ligand poses. Glide redocking was performed on structures within 30.0 kcal/mol of the best structure, but keeping only the top 20 structures overall, using XP precision and descriptors were written out.

The System Builder task was launched to generate a solvent boundary for simulation. The method was as follows: solvent model = simple point charge (SPC), box shape = orthorhombic, box size = buffer, 10 Å × 10 Å × 10 Å, exclude ion and salt placement within 20 Å of ligand, neutralize model by adding Na^+^ or Cl^−^ ions, and add 0.15 M NaCl. The MD task was launched, and the solvated system was simulated for 500 ns using Desmond. The method was as follows: simulation time = 500 ns, recording interval = 50.0 ps, ensemble class = NPT, temperature = 310.15 K, pressure = 1.01325 bar. Desmond MD jobs were processed on the Duke Compute Cluster.

For each of the prodrug-protein complexes, relative Δ*G*_binding_ values were computed on every 10^th^ frame of the final 300 ns (netting 601 frames) from the Desmond trajectories using Prime MM-GBSA.

The advanced options for Desmond MD jobs were left as default and are as follows: Integration – RESPA integrator time step: bonded = 2.0 fs, near = 2.00, far = 6.00; Ensemble – thermostat method: Nose-Hoover chain, relaxation time = 1.0 ps; barostat method: Martyna-Tobias Klein, relaxation time = 2.0 ps, coupling style – isotopic; Coulombic interaction – short range method = cutoff, cutoff radius = 9.0 Å. The relaxation process for the NPT ensemble was left as default. Details of the relaxation process are available in the Desmond user manual: (i) minimize with solute restrained; (ii) minimize without restraints; (iii) simulate in the NVT ensemble using a Berendsen thermostat with a simulation time of 12 ps, temperature of 10 K, a fast temperature relaxation constant, velocity resampling every 1 ps, nonhydrogen solute atoms restrained; (iv) simulate in the NPT ensemble using a Berendsen thermostat and Berendsen barostat with a simulation time of 12 ps, temperature of 10 K and a pressure of 1 atm, a fast temperature relaxation constant, a slow pressure relaxation constant, velocity resampling every 1 ps, nonhydrogen solute atoms restrained; (v) simulate in the NPT ensemble using a Berendsen thermostat and Berendsen barostat with a simulation time of 24 ps, temperature of 300 K and a pressure of 1 atm, a fast temperature relaxation constant, a slow pressure relaxation constant, velocity resampling every 1 ps, nonhydrogen solute atoms restrained; (vi) simulate in the NPT ensemble using a Berendsen thermostat and Berendsen barostat with a simulation time of 24 ps, temperature of 300 K and a pressure of 1 atm, a fast temperature relaxation constant, and a normal pressure relaxation constant.

### Whole Plasmid Sequencing

Transformed *E. coli* were grown overnight at 37 °C on Luria Bertani Broth (Lennox) (LB) agar plates containing the appropriate antibiotics (50 μg/mL kanamycin and/or 100 μg/mL ampicillin). A single colony was used to inoculate 5 mL of liquid LB with the appropriate antibiotics and grown at 37 °C and 200 rpm for 20 h. Cells were harvested by centrifugation (4000 rcf, 5 min, rt), resuspended in 0.5 mL of ultrapure water, and DNA was isolated using Zymo Research Zyppy Plasmid Miniprep Kit. Purified DNA was sent to plasmidsaurus for whole-plasmid Oxford Nanopore sequencing. Plasmid maps are available as Supplementary Data.

### Monoclonal Microplate Microdilution Assays

Bacteria from frozen glycerol stocks streaked onto LB agar plates containing the appropriate antibiotics (50 µg/mL kanamycin and/or 100 µg/mL ampicillin) were grown at 37 °C for 20 h. With a single colony from the agar plate, a 2 mL culture of bacteria in MHB 2 media with the appropriate antibiotics was grown at 37 °C and 200 rpm for 20 h. Prodrug stock solutions (100 mM in DMSO) were diluted to 4x of the highest tested concentration with MHB 2 media. A final solution volume of 200 µL was used for all wells in a 96-well microplate. In a 96-well microplate, 100 µL 2x serial microdilutions were performed across the long axis of the plate. Bacteria from the 2 mL culture were diluted 1:500 in 10 mL of MHB 2 media and 100 µL was added to the appropriate wells in the prepared 96-well microplate (1:1000 bacteria dilution total). The 96 well plate was incubated at 37 °C and 200 rpm for 20 h. Each microplate had a media dam around the outside of the plate, an entire row of no treatment growth control, and a positive control 2x serial microdilution of pyrithione (20 mM in DMSO), prepared as described for the prodrug microdilutions. Using a Perkin Elmer Victor^3^ V 1420 plate reader, OD_600_ was read at the initial time of plating in the microplate and subtracted from the OD_600_ readings after 20 h incubation. These values were normalized by division to the no treatment growth control for their given column.

### Bactericidal Assay

Bacteria from frozen glycerol stocks streaked onto LB agar plates containing the appropriate antibiotics (50 µg/mL kanamycin and/or 100 µg/mL ampicillin) were grown at 37 °C for 20 h. With a single colony from the agar plate, a 2 mL culture of bacteria in LB media with the appropriate antibiotics was grown at 37 °C and 200 rpm for 18 h. Overnight cultures were equalized to OD_600_ 1.0 and diluted 1000-fold in a 96-well plate filled with liquid media containing a final concentration of either 100 µM (12.7 µg/mL) pyrithione, 250 µM (31.8 µg/mL) pyrithione, 100 µM (37.5 µg/mL) AcephPT, 250 µM (94.0 µg/mL) AcephPT, or pure liquid media. The 96-well plate was incubated at 37 °C and 200 rpm for 20 h. After incubation, the contents of each of the wells were 10x serially diluted and drip-plated on LB agar plates. After 16 h of incubation at 37 °C, colony counts were enumerated to determine viable bacteria in CFU/mL.

### Profiling Hydrolysis of Prodrugs in Whole Cell Bacteria by NMR Spectroscopy

Bla-non-producing or Bla-producing engineered *E. coli* K-12 MG1655 from frozen glycerol stocks streaked onto LB agar plates containing the appropriate antibiotics (50 μg/mL kanamycin and/or 100 μg/mL ampicillin) were grown overnight at 37 °C for 20 h. With a single colony from the agar plate, a 5 mL culture of the bacteria in LB with the appropriate antibiotics was grown at 37 °C and 200 rpm for 20 h. The 5 mL culture was poured into 125 mL of LB media with the appropriate antibiotics and grown at 37 °C and 200 rpm to OD_600_ of 0.1–0.2. The cells were harvested by centrifugation (2,500 rcf, 20 min, 4 °C) and washed three times with 50 mM phosphate buffer at pH 7.5. The cells were taken up with 3 mL of 50 mM phosphate buffer in 9:1 H_2_O:D_2_O at pH/pD 7.5. Cell density was determined by diluting a sample of the cells to OD_600_ of 0.2–0.6. The cell suspension was normalized to 2.5 OD_600_ and loaded into an NMR tube. NMR experiments were taken on a Bruker Avance III 700 MHz spectrometer or a Bruker Avance III 600 MHz spectrometer. Sample temperature was set to 310 K and the sample was allowed to equilibrate before performing an experiment. A standard ^1^H NMR experiment was performed on 50 mM phosphate buffer in 9:1 H_2_O:D_2_O at pH 7.5 to determine the location of the water peak in the spectrum. A presaturation ^1^H NMR experiment was performed on the buffer sample, presaturating each scan at the determined resonance of the water peak. The cell suspension was loaded into the spectrometer and an identical presaturation ^1^H NMR experiment was performed after lock, tune, shim, and gain parameters were adjusted to collect a baseline spectrum of the cell suspension. Quickly, the cell suspension was ejected from the instrument, an aliquot of 100 mg/mL AcephPT was added to the sample by mixing with a 9” glass Pasteur pipette, and the sample was added back to the instrument. Sequential presaturation ^1^H NMR experiments were performed for the indicated time. The first scan or two were excluded from the data sets as the sample equilibrated back to 310 K. NMR spectra were processed using a Bruker TopSpin academic license. Speciation data were computed using serial integration on TopSpin. The sample containing AcephPT in buffer (*t*_0_) was used as an external calibrant to globally scale indicated resonances of sequentially acquired spectra of AcephPT incubated with *E. coli*. Resonance integration values were exported and plotted in Microsoft Excel.

### Communal Growth Competition Assay

ESBL-producing *E. coli* (16096 NDM-1-producing strain) and Bla-non-producing *E. coli* (Top10 carrying mCherry-expression plasmid) were incubated in LB with the appropriate antibiotics (50 µg/mL kanamycin and/or 100 µg/mL carbenicillin) at 37 °C and 200 rpm for 16 h. Three different overnight cultures were used as biological replicates. Overnight cultures were equalized to OD_600_ 1.0 and mixed in different ratios to 1000 µL. The mixtures were diluted 100-fold in a 96-well plate filled with liquid media containing a final concentration of either 2.2 µM (1 µg/mL) cefotaxime, 100 µM (37.5 µg/mL) AcephPT, 250 µM (94.0 µg/mL) AcephPT, or pure liquid media. Monoculture wells were also prepared as controls for each treatment. Liquid media for OD_600_ and Ex_586_/Em_620_ curves was LB, while for colony enumeration it was MHB 2. The 96-well plate was incubated at 37 °C for 24 h in a Tecan Infinite® 200Pro plate reader. Five seconds of orbital shaking followed by OD_600_ and Ex_586_/Em_620_ measurements were acquired in 10 min intervals. After incubation, the contents of each of the wells were 10x serially diluted, and drip-plated on LB agar plates containing 40 µg/mL X-gal and 1 mM IPTG. After 20 h, blue-white colony counts were enumerated to differentiate ESBL-producing *E. coli* (blue colonies) from Bla-non-producing *E. coli* (white colonies). CFU/mL was determined, and the ESBL-producer fraction was calculated by dividing the blue CFU/mL by the total CFU/mL.

### Statistical Analyses

IBM SPSS Statistics was used for all statistical analyses. For pairwise comparisons, complete datasets were assessed for significance (p<0.05) by global analysis of variance (ANOVA) before subdivision of data. If needed, additional ANOVA tests were performed before further subdivision of data and pair-wise comparison by Tukey’s group mean comparison post hoc test. For nonlinear regression, a Levenberg-Marquardt estimation method was used with a sum-of-squares convergence of 1×10^-8^ and parameter convergence of 1×10^-8^. Datasets were plotted in Microsoft Excel or RStudio.

## Abbreviations

Bla: β-lactamase
ESBL: extended-spectrum-β-lactamase
SBL: serine-β-lactamase
MBL: metallo-β-lactamase
PBP: penicillin-binding protein
RMSD: root mean squared deviation
RMSF: root mean squared fluctuation
EC_50_: half maximal effective concentration
PT: pyrithione
ARLG: Antibiotic Resistance Leadership Group
DICON: Duke Infection Control Outreach Group
DiRTE: Disinfection, Resistance, and Transmission Epidemiology Lab
NMR: nuclear magnetic resonance
DEPT: distortionless enhancement by polarization transfer
HPLC/MS: high performance liquid chromatography/mass spectrometry
HIC: hydrophobic interaction chromatography
HILIC: hydrophilic interaction chromatography
HRMS: high resolution mass spectrometry
ESI: electrospray ionization
OD_600_: optical density at 600 nm
Ex_586_/Em_620_: excitation at 585 nm, emission at 620 nm
CFU: colony forming unit
X-Gal: 5-bromo-4-chloro-3-indolyl-beta-D-galacto-pyranoside
IPTG: isopropyl β-D-1-thiogalactopyranoside
TLC: thin layer chromatography
MD: molecular dynamics
LB: Luria Bertani Broth (Lennox)
MHB: 2 Mueller Hinton Broth 2
DMSO: dimethyl sulfoxide
DCM: dichloromethane
TFA: trifluoracetic acid
TIPS: triisopropylsilane

## Acknowledgements

We thank the Duke Center for Antimicrobial Stewardship and Infection Prevention, Infection Control Outreach Network, and Antibacterial Resistance Leadership Group for maintaining and providing access to bacteria clinical isolates. We thank Dr. P. Silinski of Duke Chemistry Shared Instrumentation Facility for his assistance with HPLC/HRMS, Duke University NMR Center staff for their maintenance of NMR instruments, and Dr. M. A. Peterson and research computing staff for assistance with the Duke University Department of Chemistry Virtual Machine and use of the Duke Compute Cluster. We are grateful for financial support of this project from the Marcil-Monahan Scholars Initiative of Trinity College of Arts and Sciences, Duke University (to K.J.F.) and the National Institutes of Health (R01 GM098642 to L.Y.). We acknowledge fellowship support for A.M.D. (Duke Pharmacological Science Training Program supported by NIH T32 GM1333352 and a Burroughs Wellcome Fellowship from Duke Chemistry), A.C.J. (NSF Graduate Research Fellowship DGE 1644868) and E.A.P. and C.M.S. (Duke Trinity College Dean’s Summer Research Fellowships). We thank Duke’s Office of Undergraduate Research Support for grants to S.A.K., E.A.P. and C.M.S.

## Ethics declarations

### Competing Interests

VGF reports the following: *Grants/research support*: Astra Zeneca; MedImmune; Merck; ContraFect, Karius, Genentech, Regeneron, Basilea. *Paid Consultant*: Astra Zeneca; GSK; Armata, Debiopharm; Genentech; Basilea, Affinergy, Janssen, ContraFect, Destiny. *Royalties:* UptoDate. *Stock Options*: ArcBio, Valanbuio. *Patent pending*; sepsis diagnostics. SM, LP, FR, RM, MR, PF, JT: No reported conflicts of interest.

### Data availability

The data that support the findings in this work are included in the article, Supplementary Information, and Supplementary Data. Synthetic procedures and characterization, including NMR and mass spectral data, are including as Supplementary Information, together with details of the communal growth model, additional details from molecular dynamics simulations, clinical isolate characterization, and supplemental figures of dose-response curves, hydrolysis profiling, and bactericidal assays. Plasmid maps for engineered strains are provided as Supplementary Data, and available upon request.

## Notes

### Competing Interest Statement

The authors have declared no competing interest.

