## Supplementary Information for "An engineered prodrug selectively suppresses β-lactam resistant bacteria in a mixed microbial setting"

### Table of Contents


| | | Page |
|---|---|---|
| A | Synthetic Schemes | 3 |
| B | Synthetic Procedures and Characterization Details | 5 |
| C | NMR Spectra of Reported Compounds | 13 |
| D | HPLC/MS of Reported Compounds | 21 |
| E | Communal Growth Model | 23 |
| F | RMSDs of PBP3 Complexes from Molecular Dynamic Simulations | 25 |
| G | RMSDs of NDM-1 Complexes from Molecular Dynamic Simulations | 27 |
| H | Pyrithione Dose-Response Curves | 30 |
| I | Clinical Isolate Characterization | 31 |
| J | Profiling the Hydrolysis of AcephPT by NDM-1-producing *E. coli* | 31 |
| K | Bactericidal Assays | 32 |

### A Synthetic Schemes

#### Scheme S1. Synthesis of PcephPT

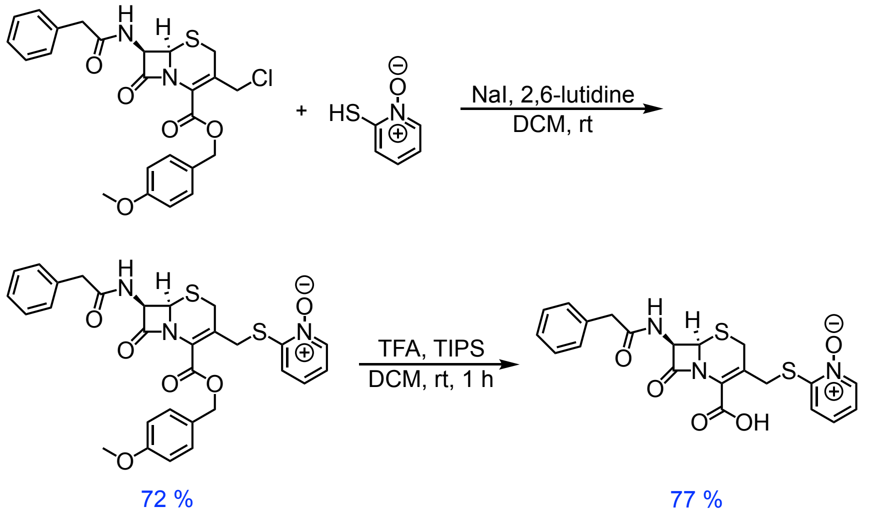


#### Scheme S2. Synthesis of AcetamidocephPT

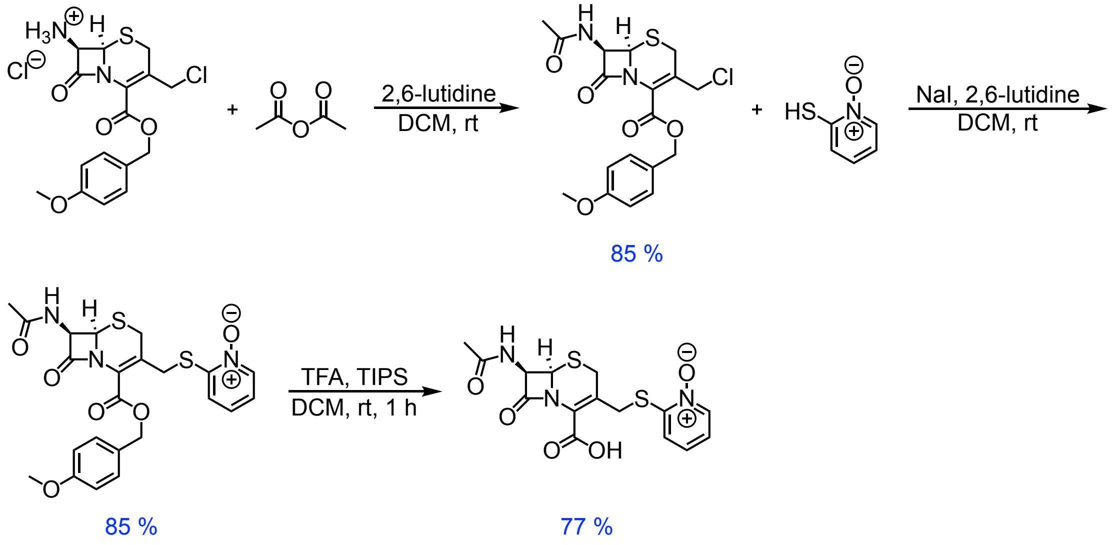

#### Scheme S3. Synthesis of AcephPT

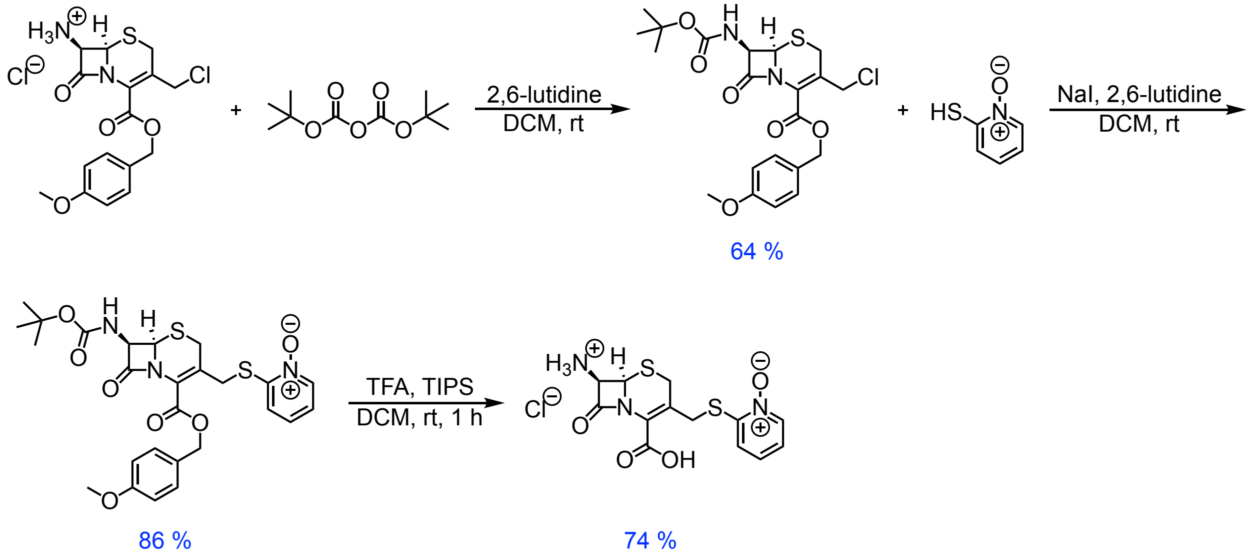

#### B Synthetic Procedures and Characterization Details

**Compound 1**. 2-((((6*R*,7*R*)-2-(((4-methoxybenzyl)oxy)carbonyl)-8-oxo-7-(2-phenylacetamido)-5-thia-1-azabicyclo[4.2.0]oct-2-en-3-yl)methyl)thio)pyridine 1-oxide

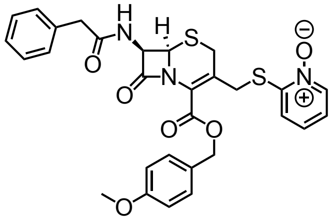

Portions of 4-methoxybenzyl (6R,7R)-3-(chloromethyl)-8-oxo-7-(2-phenylacetamido)-5-thia-1-azabicyclo[4.2.0]oct-2-ene-2-carboxylate (1.4610 g; 3.00 mmol), sodium iodide (0.4507 g; 3.01 mmol), and 25 mL of dichloromethane were added to an oven-dried 100 mL round-bottom flask. After stirring for 1 h, 2-mercaptopyridine *N*-oxide (0.4578 g, 3.60 mmol) and 2,6-lutidine (0.700 mL, 6.04 mmol) were added and the mixture was stirred for an additional 4 h followed by separation into a separatory funnel with 50 mL of dichloromethane and 50 mL of DI-water. After agitation, the organic layer was separated and the aqueous was extracted with 2 x 50 mL of dichloromethane. Organics were combined, washed 3 x 50 mL of 1 M HCl, 50 mL of DI-water, and 50 mL of brine. Organics were dried over anhydrous $MgSO_4$ and solvents were removed by rotary evaporation. Purification was performed by column chromatography, using 23:2 dichloromethane:methanol as an eluent. The purified product was dried on high vacuum overnight to provide 1.2474 g (72 % yield) of a light grey solid.

$^1$H NMR: (500 MHz, $CDCl_3$): δ = 8.22 (d, *J* = 6.3 Hz, 1H), 7.36-7.24 (m, 7H), 7.22 (dd, *J* = 1.6, 8.2 Hz, 1H), 7.16 (t, *J* = 7.7 Hz, 1H), 7.10 (dt, *J* = 1.7, 10.4 Hz, 1H), 7.05 (d, *J* = 9.0 Hz, 1H), 6.87 (d, *J* = 8.6 Hz, 2H), 5.79 (dd, *J* = 4.8, 9.0 Hz. 1H), 5.22 (s, 2H), 4.88 (d, *J* = 4.9 Hz, 1H), 4.10 (d, *J* = 12.8 Hz, 1H), 4.02 (d, *J* = 12.8 Hz, 1H), 3.81 (s, 3H), 3.67 (s, 2H), 3.60-3.51 (ABq, 2H).

$^{13}$C{$^1$H} NMR: (125 MHz, $CDCl_3$): δ = 171.52, 164.91, 161.66, 160.03, 150.43, 139.12, 134.21, 130.83, 129.46, 129.07, 127.56, 126.76, 126.54, 126.25, 125.67, 123.24, 121.65, 114.08, 68.19, 59.40, 58.00, 55.36, 43.23, 33.03, 28.07.

HRMS (ESI): *m/z* $[M + H]^+$ calculated for $[C_{29}H_{27}N_3O_6S_2 + H]^+$: 578.1414; found: 578.1431

**PcephPT**. 2-((((6*R*,7*R*)-2-carboxy-8-oxo-7-(2-phenylacetamido)-5-thia-1-azabicyclo[4.2.0]oct-2-en-3-yl)methyl)thio)pyridine 1-oxide

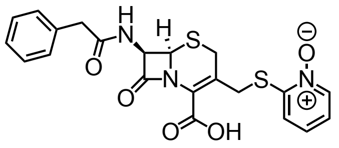

Compound 1 (0.5779 g, 1.00 mmol) was added to an oven-dried 100 mL round-bottom flask. A cocktail of 12 mL dichloromethane, 7 mL of trifluoroacetic acid, and 1 mL of triisopropylsilane was prepared in a beaker and added to the flask. After 60 min, solvent was removed by rotary evaporation. The residue was dissolved in 5 mL of methanol and added dropwise to 300 mL of rapidly stirring diethyl ether. The suspension was stirred for 15 min before the precipitate was collected by vacuum filtration. The product was dried under high vacuum overnight to provide 0.3526 g (77 %) of a white solid.

$^1$H NMR: (500 MHz, $d_6$-DMSO): δ = 9.13 (d, $J$ = 8.4 Hz, 1H), 8.31 (d, $J$ = 6.3 Hz, 1H), 7.48 (dd, $J$ = 1.0, 8.2 Hz, 1H), 7.35 (t, $J$ = 7.6 Hz, 1H), 7.32-7.26 (m, 4H), 7.25-7.20 (m, 2H), 5.67 (dd, $J$ = 4.8, 8.3, 1H), 5.12 (d, $J$ = 4.8 Hz, 1H), 4.12-4.03 (ABq, 2H), 3.74 (d, $J$ = 18.0 Hz, 1H), 3.57 (d, $J$ = 12.6 Hz, 1H), 3.56 (d, $J$ = 18.3 Hz, 1H), 3.49 (d, $J$ = 13.9 Hz, 1H).

$^{13}$C{$^1$H} NMR: (125 MHz, $d_6$-DMSO): δ = 171.13, 164.77, 163.09, 150.25, 138.41, 135.86, 129.11, 128.34, 126.61, 126.31, 125.82, 125.13, 122.33, 121.62, 59.15, 57.87, 41.70, 32.40, 27.41.

HRMS (ESI): $m/z$ $[M + H]^+$ calculated for $[C_{21}H_{19}N_3O_5S_2 + H]^+$: 458.0839; found: 458.0839

**Compound 2**. 4-methoxybenzyl (6*R*,7*R*)-7-acetamido-3-(chloromethyl)-8-oxo-5-thia-1-azabicyclo[4.2.0]oct-2-ene-2-carboxylate

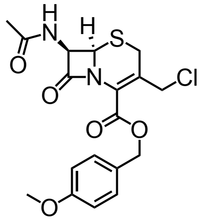

Portions of 4-methoxybenzyl (6*R*,7*R*)-7-amino-3-(chloromethyl)-8-oxo-5-thia-1-azabicyclo[4.2.0]oct-2-ene-2-carboxylate hydrochloride (1.6219 g; 4.00 mmol) was dissolved in 100 mL dichloromethane in an oven-dried 250 mL round-bottom flask. Acetic anhydride (1.550 mL, 16.40 mmol) and 2,6-lutidine (0.930 mL, 8.03 mmol) were added to the mixture. After 2 h, the mixture was poured into a separatory funnel with 100 mL of brine and 100 mL ethyl acetate. After agitation, the organic layer was collected and the aqueous layer was extracted with 2 x 100 mL of ethyl acetate. Organics were combined and dried over anhydrous $MgSO_4$. Solvent was removed by rotary evaporation. The product was dried on high vacuum overnight to provide 1.4366 g (85 % yield) of white solid.

$^1$H NMR: (500 MHz, $CDCl_3$): δ = 7.34 (d, *J* = 8.7 Hz, 2H), 6.89 (d, *J* = 8.7, 2H), 6.29 (d, *J* = 9.0, 1H), 5.84 (dd, *J* = 5.0, 9.0 Hz, 1H), 5.26-5.18 (ABq, 2H), 4.96 (d, *J* = 5.0 Hz, 1H), 4.53 (d, *J* = 11.8 Hz, 1H), 4.43 (d, *J* = 11.8 Hz, 1H), 3.81 (s, 3H), 3.65 (d, *J* = 18.3 Hz, 1H), 3.48 (d, *J* = 18.3 Hz, 1H), 2.06 (s, 3H).

$^{13}$C{$^1$H} NMR: (125 MHz, $CDCl_3$): δ = 170.31, 165.15, 161.26, 160.13, 130.84, 126.73, 126.33, 125.76, 114.16, 68.40, 59.43, 57.77, 55.42, 43.40, 27.32, 22.98.

HRMS (ESI): *m*/*z* $[M - H]^-$ calculated for $[C_{18}H_{19}ClN_2O_5S - H]^-$: 409.0625; found: 409.0627

**Compound 3**. 2-((((6*R*,7*R*)-7-acetamido-2-(((4-methoxybenzyl)oxy)carbonyl)-8-oxo-5-thia-1-azabicyclo[4.2.0]oct-2-en-3-yl)methyl)thio)pyridine 1-oxide

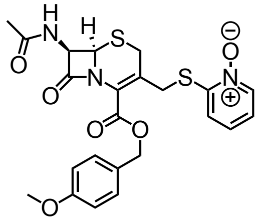

Compound 2 (1.2332 g; 3.00 mmol), sodium iodide (0.4596 g; 3.07 mmol), and 25 mL of dichloromethane were added to an oven-dried 100 mL round-bottom flask and stirred for 1 h. 2-mercaptopyridine *N*-oxide (0.4596 g, 3.59 mmol) and 2,6-lutudine (0.700 mL, 6.04 mmol) were added to the mixture. After an additional 4 h, the mixture was poured into a separatory funnel with 50 mL of dichloromethane and 50 mL of DI-water. After agitation, the organic layer was separated and the aqueous was extracted with 2 x 50 mL of dichloromethane. Organics were combined, washed 3 x 50 mL of 1 M HCl, 50 mL of DI-water, and 50 mL of brine. Organics were dried over anhydrous $MgSO_4$ and solvents were removed by rotary evaporation. Purification was performed by column chromatography, using 23:2 dichloromethane:methanol as an eluent. The purified product was dried on high vacuum overnight to provide 1.2836 g (85 % yield) of a white solid.

$^{1}$H NMR: (500 MHz, $CDCl_3$): δ = 8.19 (d, *J* = 6.4 Hz, 1H), 7.36-7.29 (m, 3H), 7.21 (dd, *J* = 1.6, 8.2 Hz, 1H), 7.16 (t, *J* = 7.63 Hz, 1H), 7.08 (dt, *J* = 1.7, 6.9 Hz, 1H), 6.84 (d, *J* = 8.65 Hz, 2H), 5.81 (dd, *J* = 5.1, 9.2 Hz, 1H), 5.20 (s, 2H), 4.90 (d, *J* = 4.9 Hz, 1H), 4.13 (d, *J* = 12.6 Hz, 1H), 3.96 (d, *J* = 12.7 Hz, 1H), 3.78 (s, 3H), 3.65-3.53 (ABq, 2H), 2.07 (s, 3H).

$^{13}$C{$^{1}$H} NMR: (125 MHz, $CDCl_3$): δ = 170.83, 165.37, 161.72, 160.04, 150.56, 139.15, 130.82, 126.79, 126.50, 126.38, 125.71, 123.27, 121.68, 114.10, 68.20, 59.47, 57.99, 55.38, 33.13, 28.13, 22.90.

HRMS (ESI): *m*/*z* $[M + H]^+$ calculated for $[C_{23}H_{23}N_3O_6S_2 + H]^+$: 502.1101; found: 502.1093

**AcetamidecephPT**. 2-((((6*R*,7*R*)-7-acetamido-2-carboxy-8-oxo-5-thia-1-azabicyclo[4.2.0]oct-2-en-3-yl)methyl)thio)pyridine 1-oxide

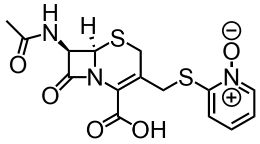

Compound 3 (0.5021 g, 1.00 mmol) was added to an oven-dried 100 mL round-bottom flask. A cocktail of 12 mL dichloromethane, 7 mL of trifluoroacetic acid, and 1 mL of triisopropylsilane was prepared in a beaker and added to the flask. After 60 min, solvent was removed by rotary evaporation. The residue was dissolved in 5 mL of methanol and added dropwise to 300 mL of rapidly stirring diethyl ether. The suspension was left to stir for 15 min before the precipitate was collected by vacuum filtration. The product was dried under high vacuum overnight to provide 0.2928 g (77 %) of a white solid.

$^{1}$H NMR: (500 MHz, $d_6$-DMSO): δ = 8.86 (d, *J* = 8.4 Hz, 1H), 8.31 (d, *J* = 5.9 Hz, 1H), 7.48 (dd, *J* = 1.6, 8.3 Hz, 1H), 7.35 (dt, *J* = 0.9, 7.9 Hz, 1H), 7.23 (dt, *J* = 1.6, 7.0 Hz, 1H), 5.67 (dd, *J* = 4.8, 8.4 Hz, 1H), 5.12 (d, *J* = 4.8 Hz, 1H), 4.12-4.03 (ABq, 2H), 3.74 (d, *J* = 18.0 Hz, 1H), 3.56 (d, *J* = 17.9 Hz, 1H), 1.91 (s, 3H).

$^{13}$C{$^{1}$H} NMR: (125 MHz, $d_6$-DMSO): δ = 170.05, 164.95, 163.03, 150.19, 138.34, 126.24, 125.65, 125.12, 122.28, 121.54, 59.03, 57.77, 32.32, 27.36, 22.07.

HRMS (ESI): *m/z* $[M + H]^+$ calculated for $[C_{15}H_{15}N_3O_5S_2 + H]^+$: 382.0526; found: 382.0516

**Compound 4**. 4-methoxybenzyl (6*R*,7*R*)-7-((*tert*-butoxycarbonyl)amino)-3-(chloromethyl)-8-oxo-5-thia-1-azabicyclo[4.2.0]oct-2-ene-2-carboxylate

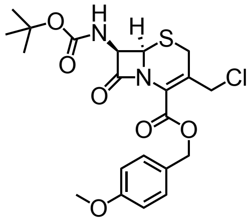

Portions of 4-methoxybenzyl (6*R*,7*R*)-7-acetamide-3-(chloromethyl)-8-oxo-5-thia-1-azabicyclo[4.2.0]oct-2-ene-2-carboxylate (2.0297 g, 5.01 mmol) and 100 mL of dichloromethane were added to an oven-dried 250 mL round-bottom flask. Di-*tert*-butyl dicarbonate (4.3649 g, 20.00 mmol) and 2,6-lutidine (1.200 mL, 10.36 mmol) were added to the mixture. After stirring at rt for 2.5 h, the mixture was poured into a separatory funnel with 100 mL of brine and 100 mL of ethyl acetate. After agitation, the organic layer was separated and the aqueous was extracted with 2 x 100 mL of ethyl acetate. Organics were combined and dried over anhydrous $MgSO_4$. Solvents were removed by rotary evaporation. Purification was performed by column chromatography, using 2:1 hexanes:ethyl acetate as an eluent. The purified product was dried under vacuum overnight to provide 1.5509 g (64 % yield) of a white solid.

$^1$H NMR: (500 MHz, $CDCl_3$): δ = 7.34 (d, *J* = 8.7 Hz, 2H), 6.89 (d, *J* = 8.7 Hz, 2H), 5.61 (dd, *J* = 4.8, 9.5 Hz, 1H), 5.27-5.20 (m, 3H), 4.94 (d, *J* = 4.9 Hz, 1H), 4.56 (d, *J* = 11.8, 1H), 4.42 (d, *J* = 11.8 Hz, 1H), 3.81 (s, 3H), 3.65 (d, *J* = 18.3 Hz, 1H), 3.47 (d, *J* = 18.7 Hz, 1H), 1.45 (s, 9H).

$^{13}$C{$^1$H} NMR: (125 MHz, $CDCl_3$): δ = 165.49, 161.33, 160.09, 154.63, 130.84, 126.77, 126.23, 125.79, 114.14, 81.41, 68.36, 61.16, 58.19, 55.40, 43.48, 28.30, 27.27.

HRMS (ESI): *m/z* $[M - H]^-$ calculated for $[C_{21}H_{25}ClN_2O_6S - H]^-$: 467.1044; found: 467.1045

**Compound 5**. 2-((((6*R*,7*R*)-7-((*tert*-butoxycarbonyl)amino)-2-(((4-methoxybenzyl)oxy) carbonyl)-8-oxo-5-thia-1-azabicyclo[4.2.0]oct-2-en-3-yl)methyl)thio)pyridine 1-oxide

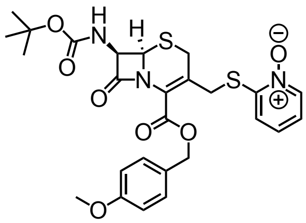

Compound 4 (1.1716 g, 2.50 mmol), sodium iodide (0.3780 g, 2.52 mmol), and 25 mL of dichloromethane were added to an oven-dried 100 mL round-bottom flask and allowed to stir for 1 h. 2-mercaptopyridine *N*-oxide (0.3820 g, 3.00 mmol) and 2,6-lutidine (0.580 mL, 5.01 mmol) were added to the mixture. After an additional 4 h, the mixture was poured into a separatory funnel with 50 mL of dichloromethane and 50 mL of DI-water. After agitation, the organic layer was separated and the aqueous was extracted with 2 x 50 mL of dichloromethane. Organics were combined, washed 3 x 50 mL of 100 mM HCl, 50 mL of DI-water, and 50 mL of brine. Organics were dried over anhydrous $MgSO_4$ and solvents were removed by rotary evaporation. Purification was performed by column chromatography, using 23:2 dichloromethane:methanol as an eluent. The purified product was dried under high vacuum overnight to provide 1.2026 g (86 % yield) of a light-purple solid.

$^1$H NMR: (500 MHz, $CDCl_3$): δ = 8.21 (d, *J* = 8.2 Hz, 1H), 7.32 (d, *J* = 8.6 Hz, 2H), 7.17 (d, *J* = 8.1 Hz, 1H), 7.11 (t, *J* = 7.6 Hz, 1H), 7.05 (d, *J* = 6.7 Hz, 1H), 6.84 (d, *J* = 8.6 Hz, 2H), 5.56 (dd, *J* = 4.4, 9.3 Hz, 1H), 5.48 (d, *J* = 10.1 Hz, 1H), 5.20 (s, 2H), 4.90 (d, *J* = 4.6 Hz, 1H), 4.17-4.03 (ABq, 2H), 3.78 (m, 3H), 3.67-3.55 (ABq, 2H), 1.43 (s, 9H).

$^{13}$C{$^1$H} NMR: (125 MHz, $CDCl_3$): δ = 165.42, 161.71, 159.97, 154.65, 150.32, 138.95, 130.79, 126.77, 126.70, 125.86, 125.69, 123.12, 121.52, 114.04, 81.17, 68.15, 61.04, 58.25, 55.32, 33.02, 28.23, 28.13.

HRMS (ESI): *m*/*z* $[M + H]^+$ calculated for $[C_{26}H_{29}N_3O_7S_2 + H]^+$: 560.1520; found: 560.1525

**AcephPT**. 2-((((6*R*,7*R*)-7-amino-2-carboxy-8-oxo-5-thia-1-azabicyclo[4.2.0]oct-2-en-3-yl)methyl)thio)pyridine 1-oxide hydrochloride

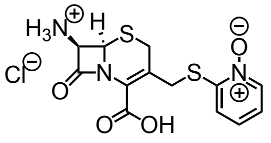

Compound 5 (0.5599 g, 1.00 mmol) was added to an oven-dried 100 mL round-bottom flask. A cocktail of 12 mL dichloromethane, 7 mL of trifluoroacetic acid, and 1 mL of triisopropylsilane was prepared in a beaker and added to the flask. After 60 min, solvent was removed by rotary evaporation. The residue was dissolved in 5 mL of 3.0 M methanolic HCl and added dropwise to 300 mL of rapidly stirring diethyl ether. The suspension was left to stir for 15 min before the precipitate was collected by vacuum filtration. The product was dried on the high vacuum overnight to provide 0.2780 g (74 %) of a white solid.

$^{1}$H NMR: (500 MHz, $d_6$-DMSO): δ = 9.36 (br, 2H), 8.32 (d, *J* = 6.3 Hz, 1H), 7.51 (dd, *J* = 1.2, 8.5 Hz, 1H), 7.39 (t, *J* = 7.8 Hz, 1H), 7.26 (t, *J* = 7.7 Hz, 1H), 5.22 (d, *J* = 4.9 Hz, 1H), 5.12 (d, *J* = 4.9 Hz, 1H), 4.22-4.14 (ABq, 2H), 3.77-3.68 (ABq, 2H).

$^{13}$C{$^{1}$H} NMR: (125 MHz, $d_6$-DMSO): δ = 162.66, 160.23, 150.22, 138.56, 131.04, 126.38, 126.19, 122.55, 121.83, 57.93, 54.66, 32.11, 28.06.

HRMS (ESI): *m/z* $[M + H]^+$ calculated for $[C_{13}H_{13}N_3O_4S_2 + H]^+$: 340.0420; found: 340.0416

#### C NMR Spectra of Reported Compounds

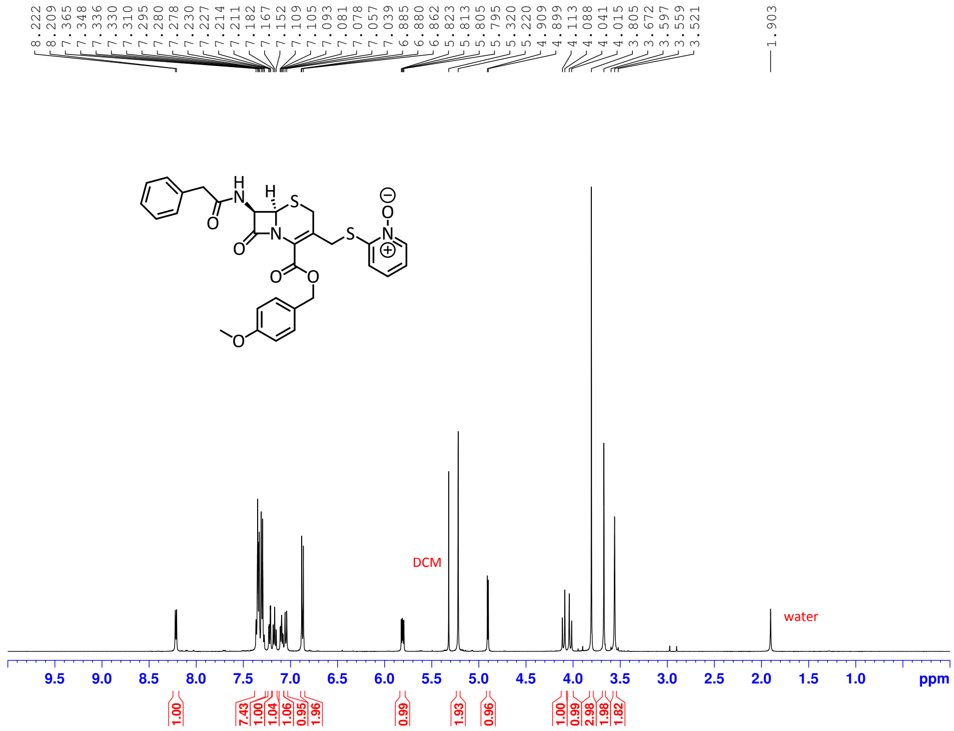


**Figure S1. $^{1}$H NMR of Compound 1**

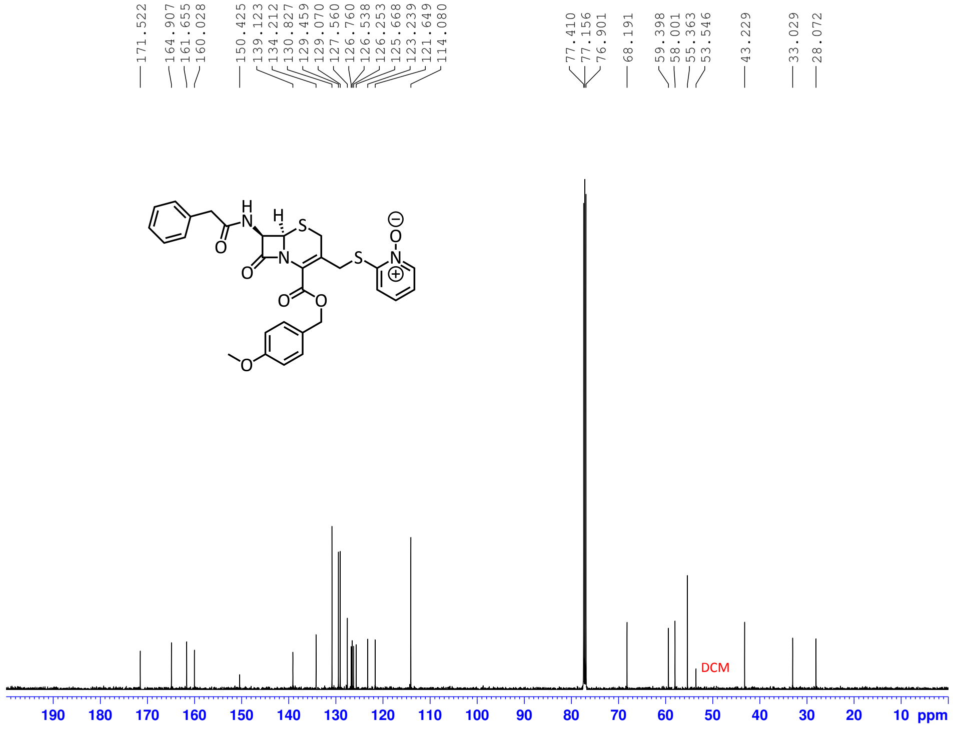


**Figure S2. $^{13}$C{$^{1}$H} NMR of Compound 1**

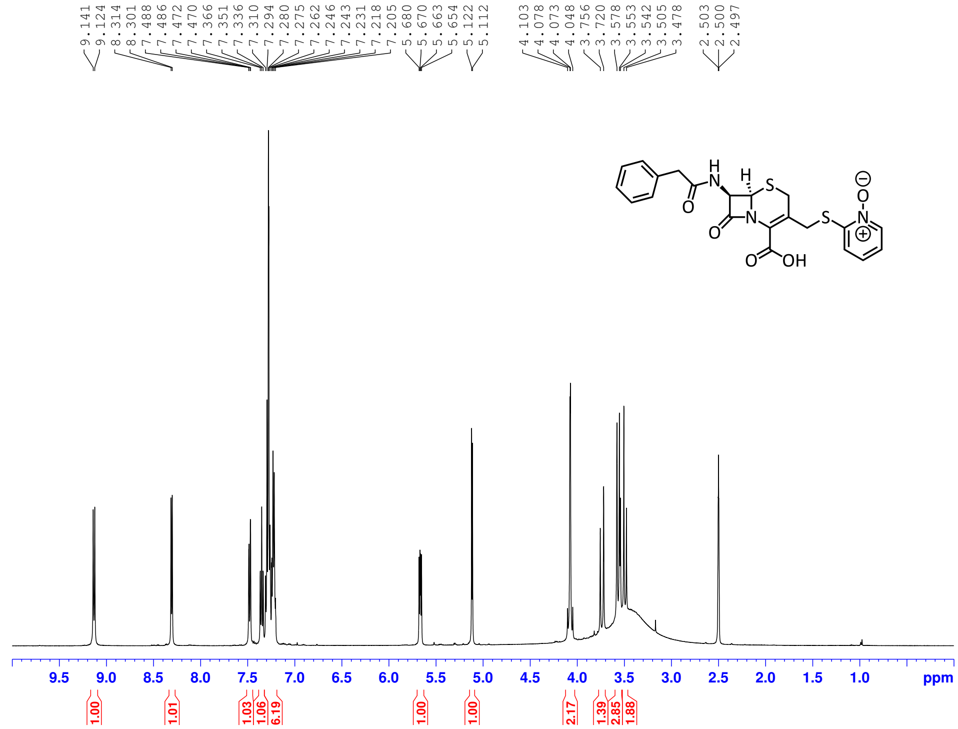


**Figure S3.** $^{1}$H **NMR of PcephPT**

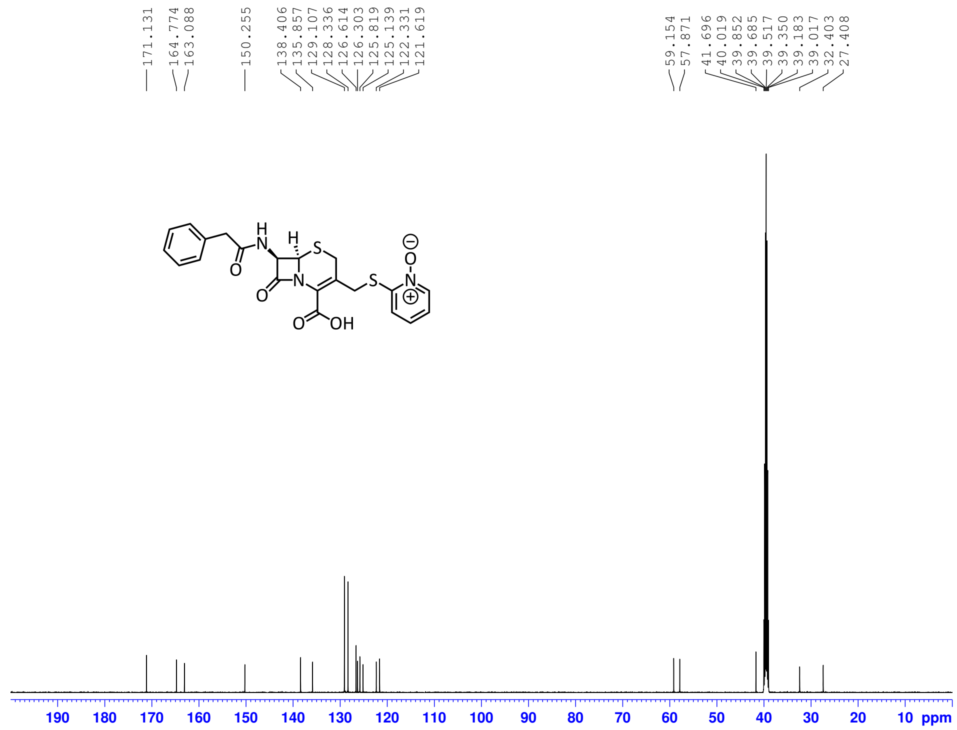


**Figure S4.** $^{13}$C{$^{1}$H} **NMR of PcephPT**

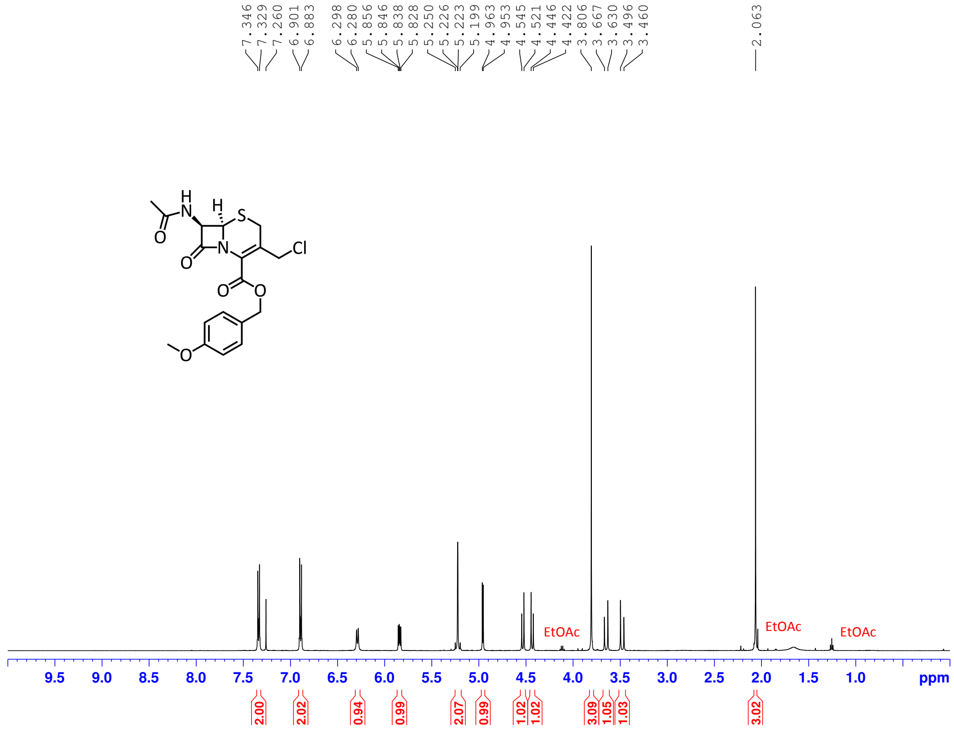


**Figure S5. ¹H NMR of Compound 2**

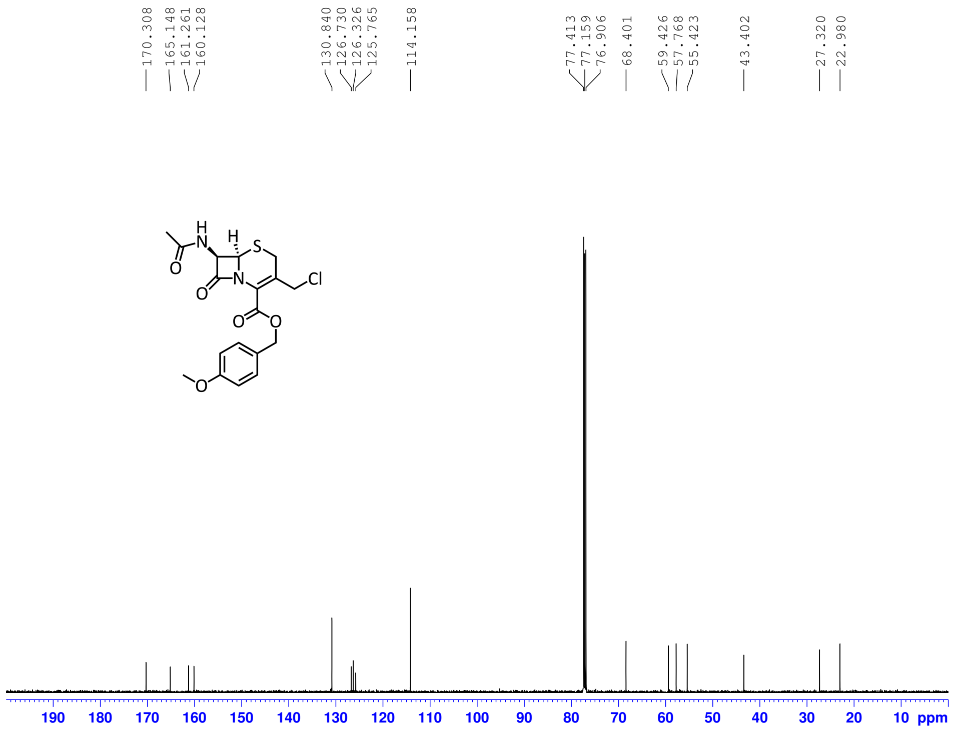


**Figure S6. $^{13}C\{^1H\}$ NMR of Compound 2**

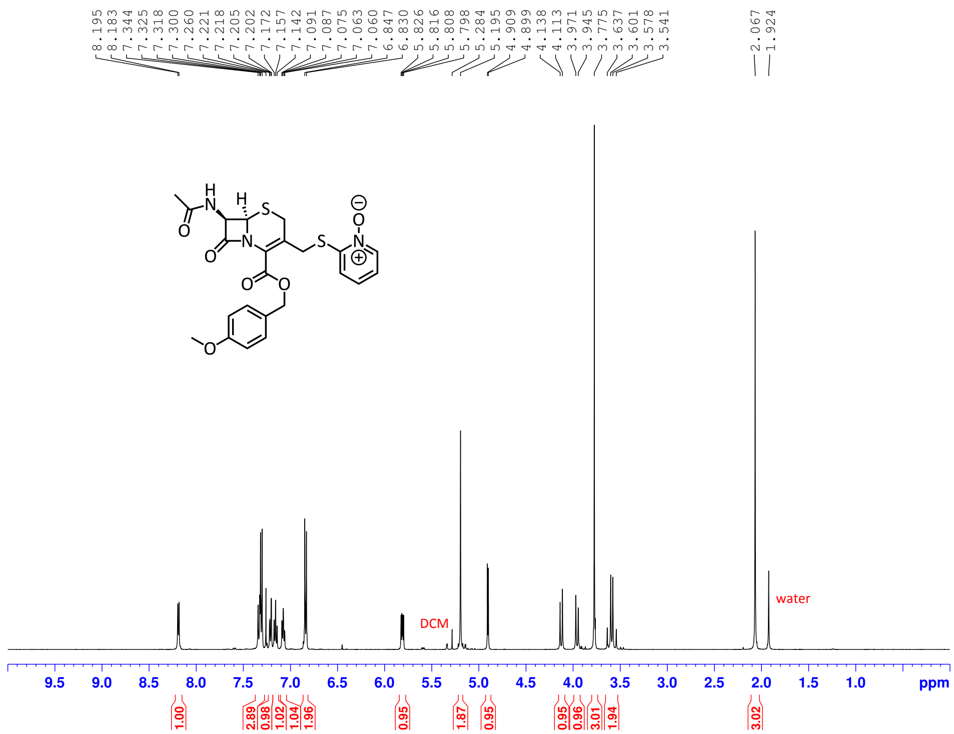


**Figure S7.** $^{1}H$ **NMR of Compound 3**

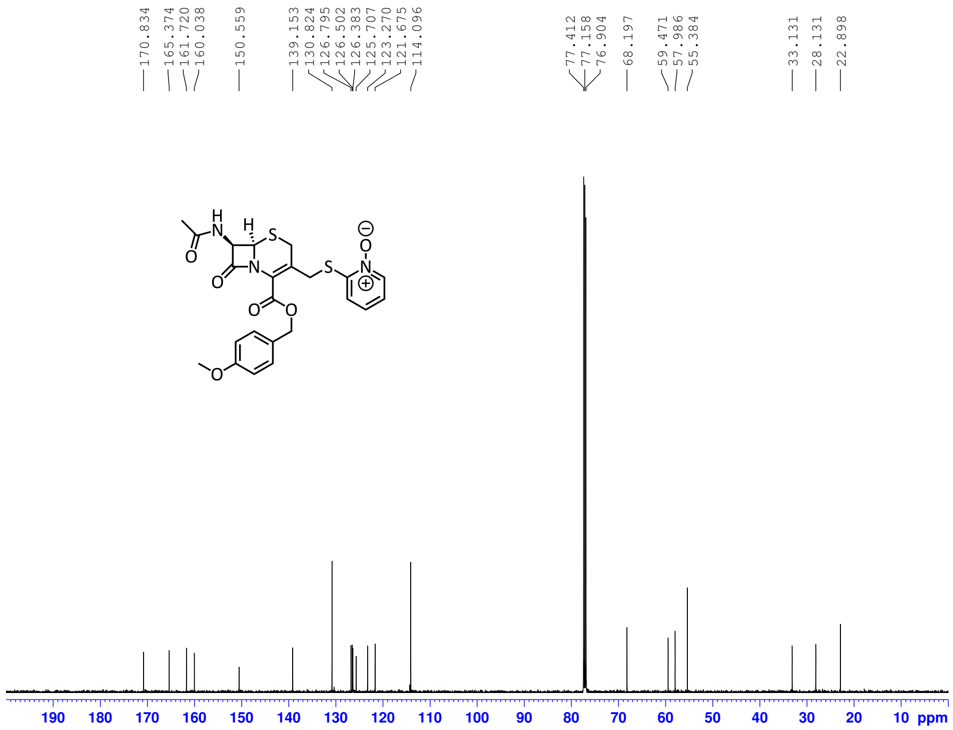


**Figure S8.** $^{13}C\{^{1}H\}$ **NMR of Compound 3**

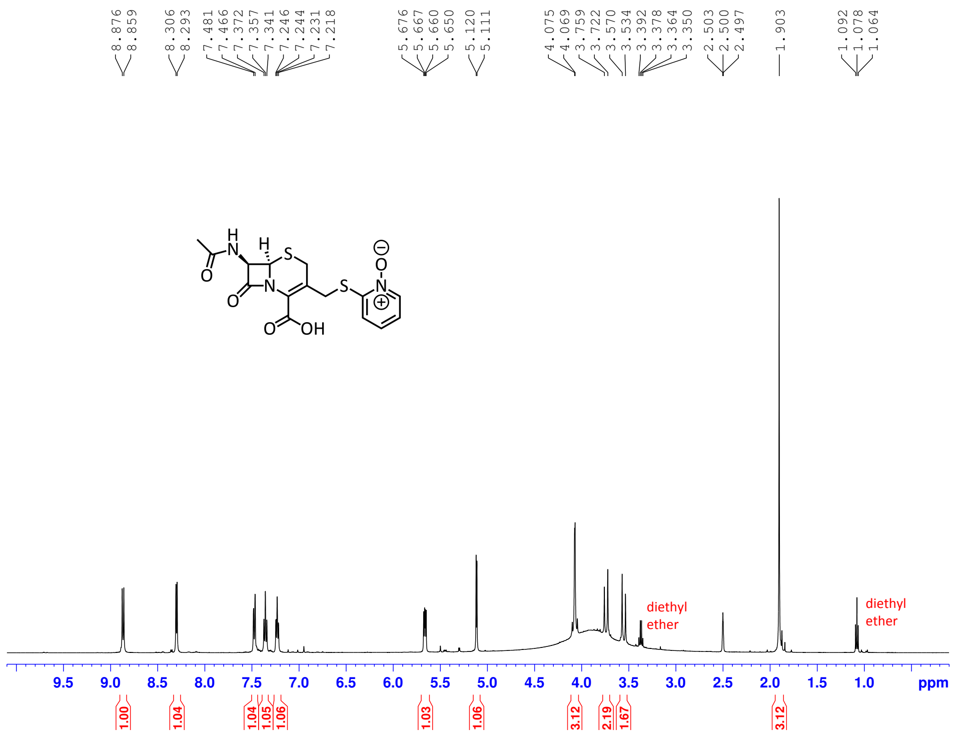


**Figure S9.** $^{1}$**H NMR of AcetamidecephPT**

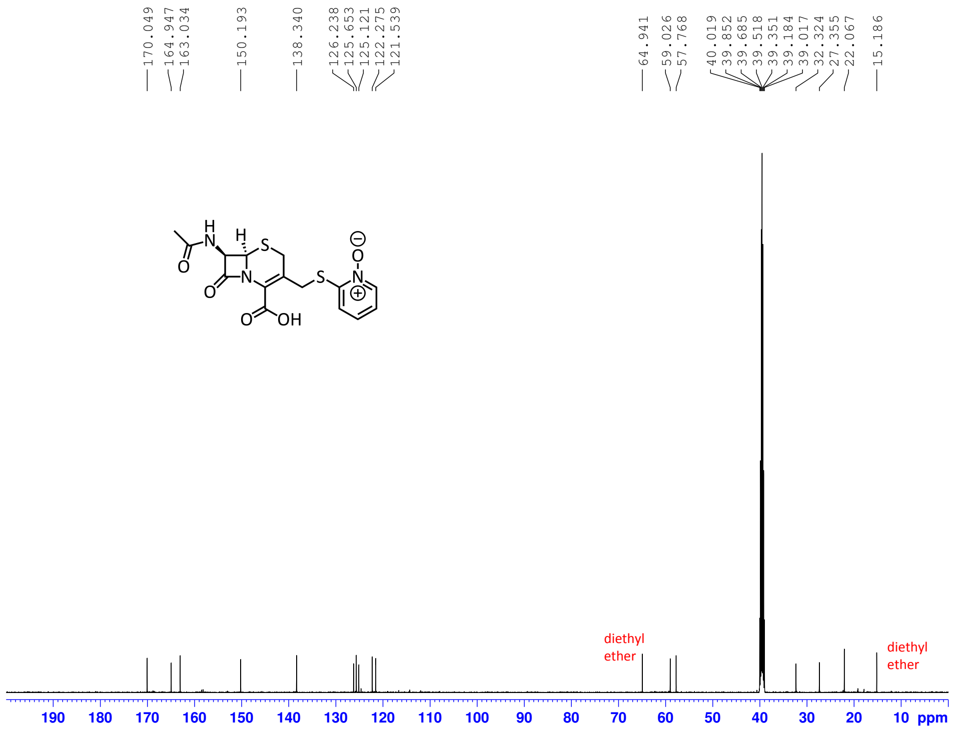


**Figure S10.** $^{13}$**C{**$^{1}$**H} NMR of AcetamidecephPT**

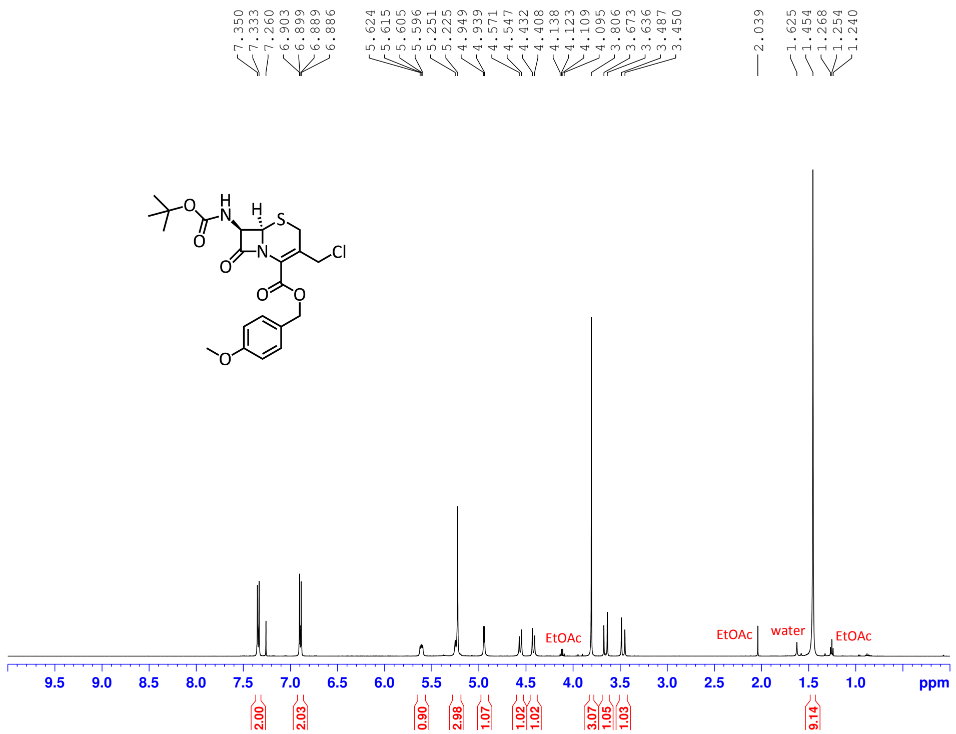


**Figure S11. [1]H NMR of Compound 4**

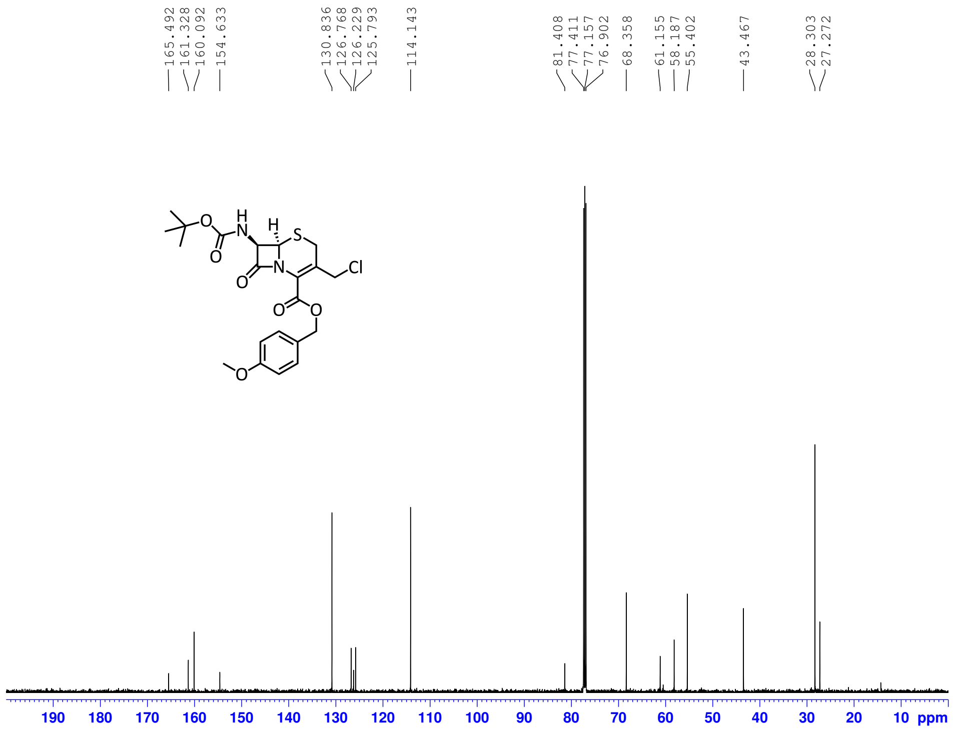


**Figure S12. [13]C{[1]H} NMR of Compound 4**

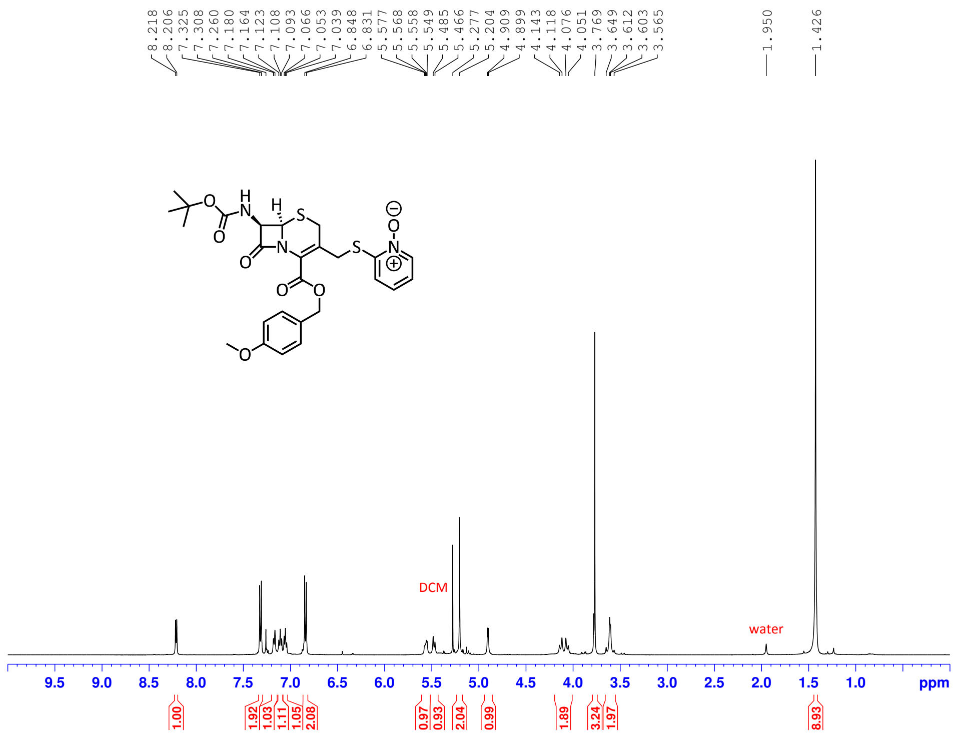


**Figure S13. $^{1}$H NMR of Compound 5**

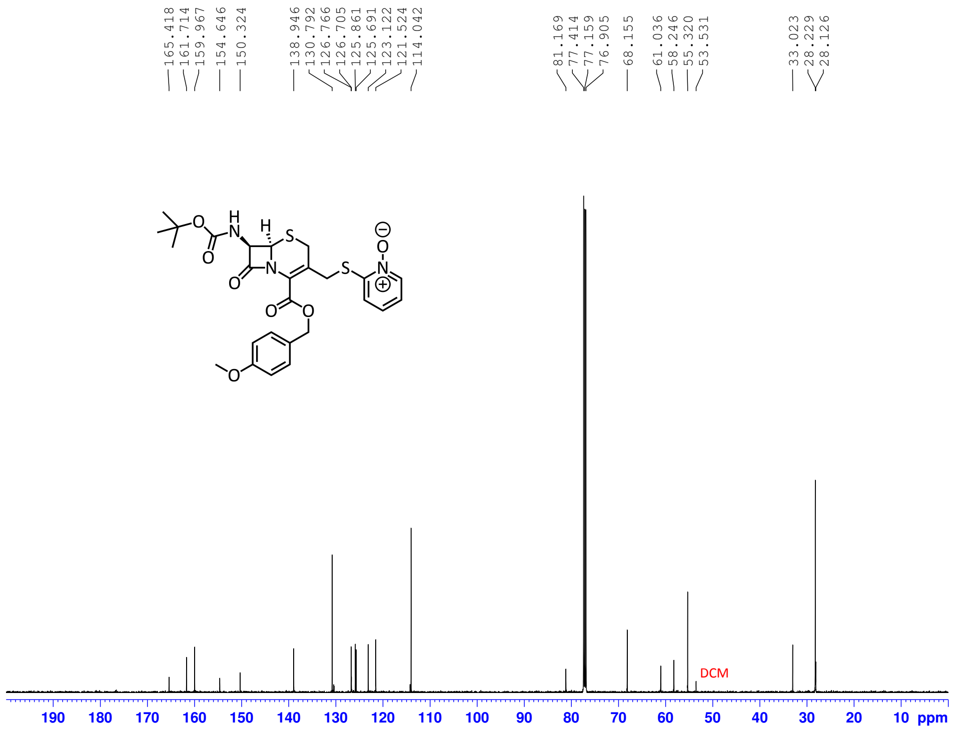


**Figure S14. $^{13}$C{$^{1}$H} NMR of Compound 5**

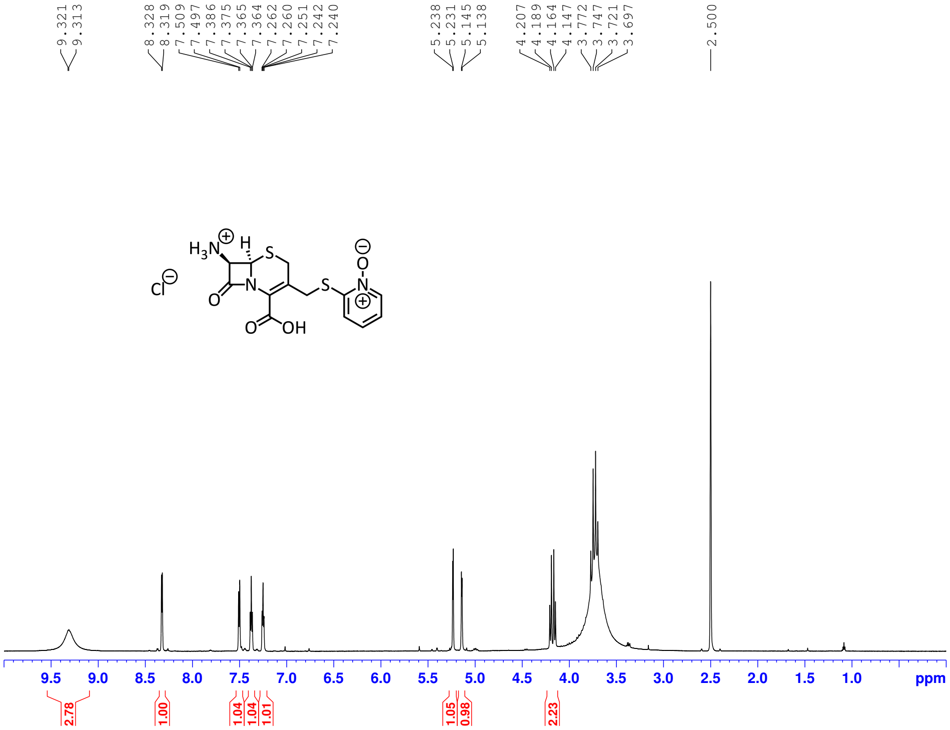


**Figure S15. ¹H NMR of AcephPT**

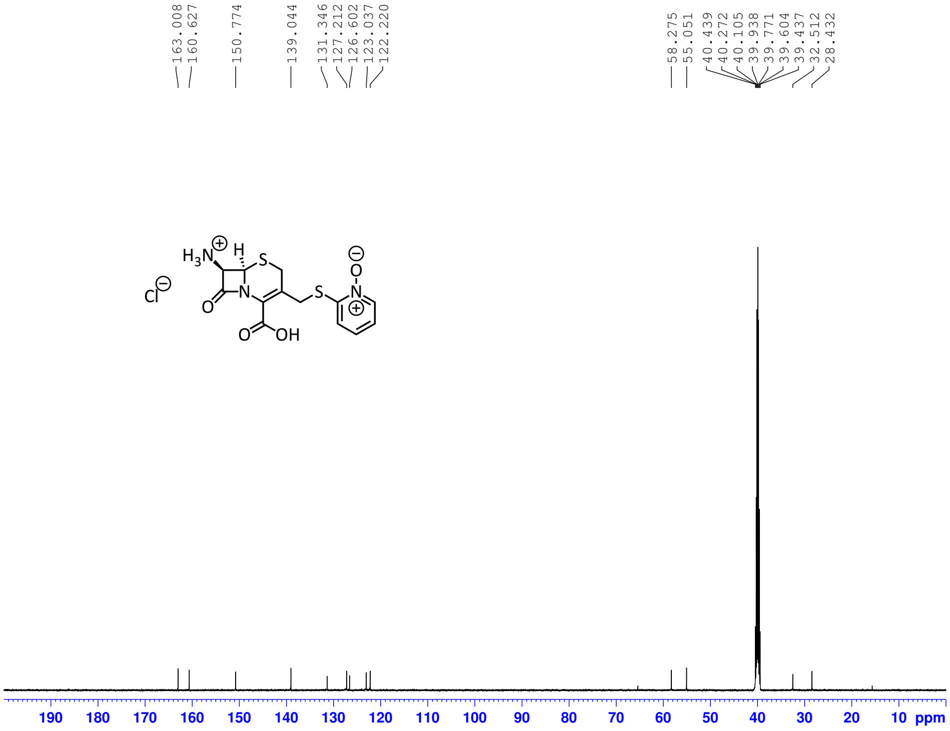


**Figure S16. $^{13}C\{^1H\}$ NMR of AcephPT**

#### D HPLC/MS of Reported Compounds

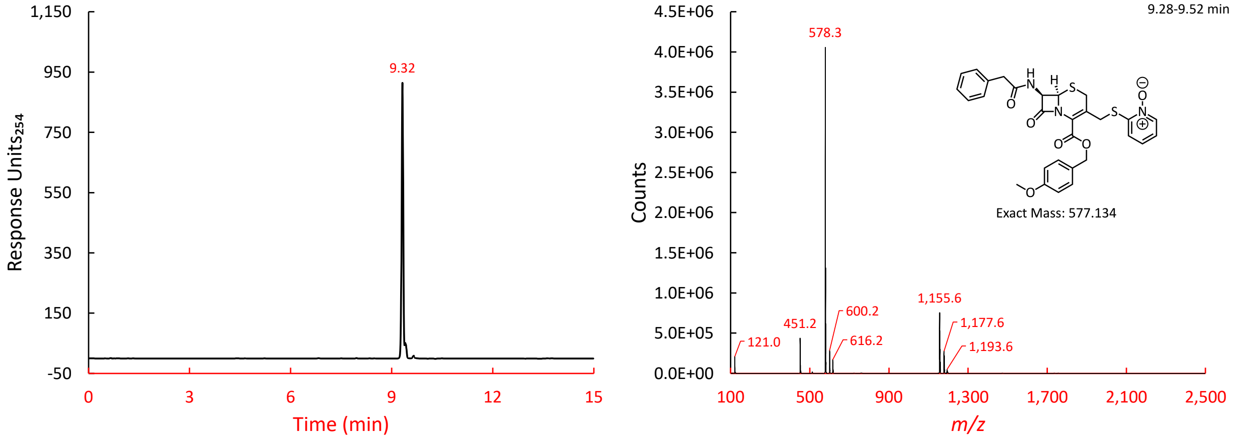


**Figure S17. HPLC/MS of Compound 1 by HIC HPLC**

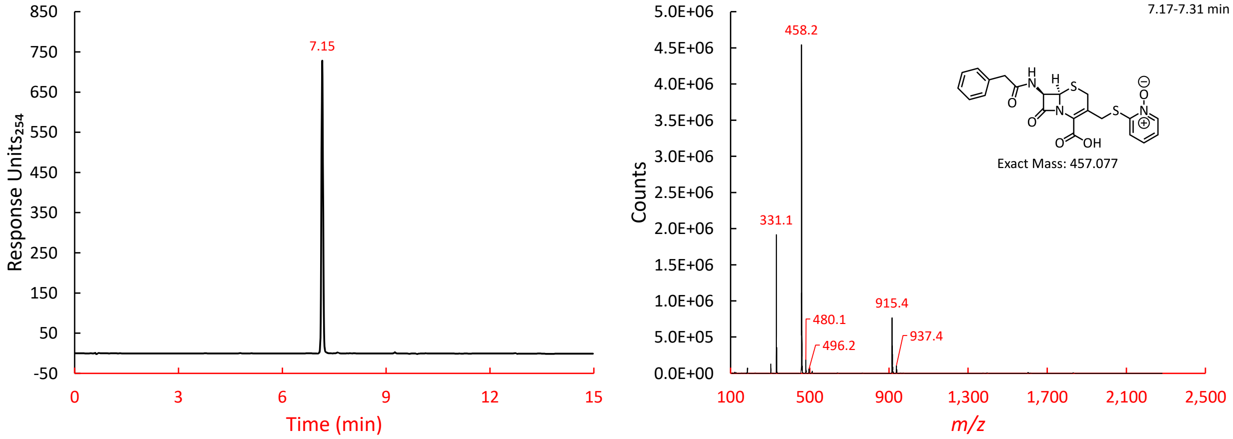


**Figure S18. HPLC/MS of PcephPT by HIC HPLC**

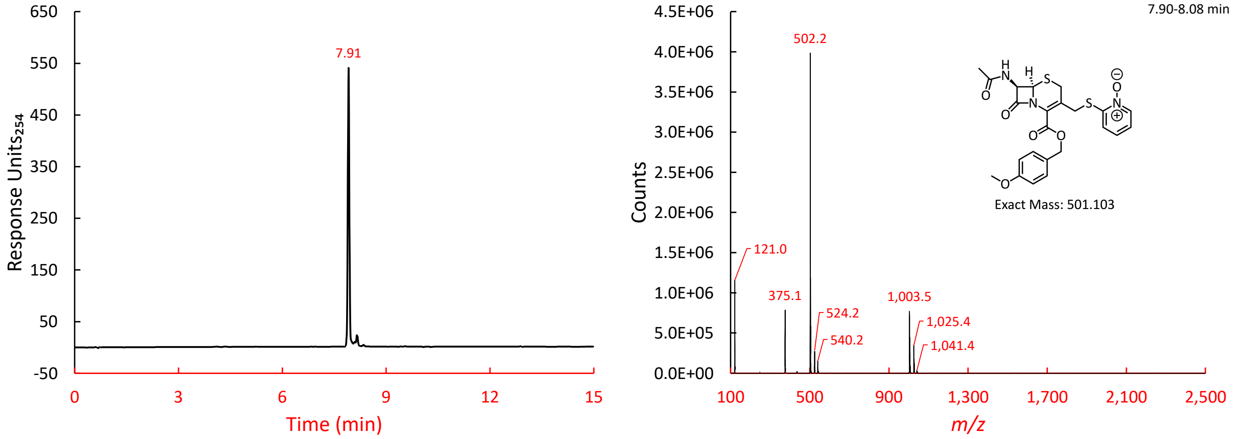


**Figure S19. HPLC/MS of Compound 3 by HIC HPLC**

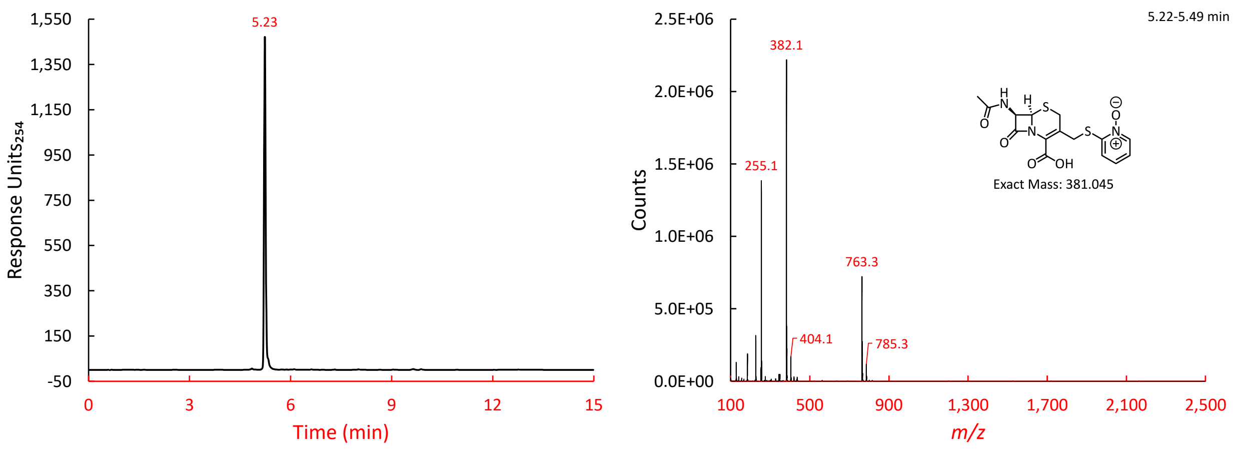


Figure S20. HPLC/MS of AcetamidecephPT by HIC HPLC

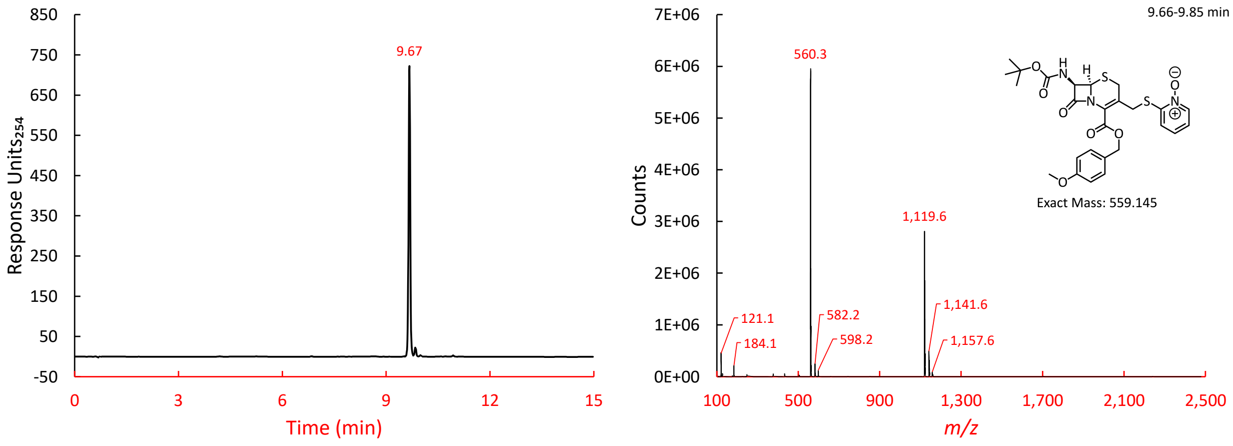


Figure S21. HPLC/MS of Compound 5 by HIC HPLC

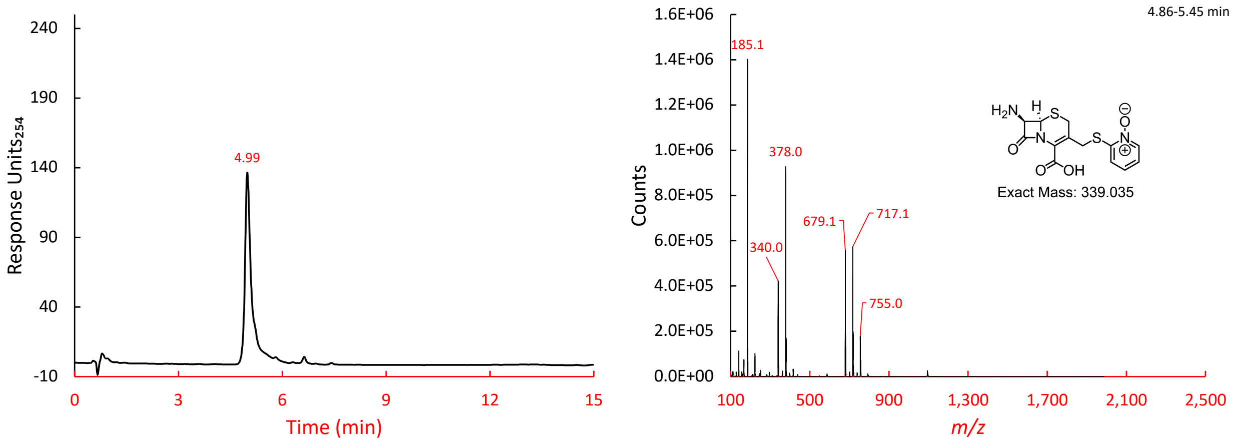


Figure S22. HPLC/MS of AcephPT by HILIC HPLC

#### E Communal Growth Model

The differential equations formulated to simulate the qualitative dynamics shown in Figure 1 are shown below in Eqs 1–10. The dynamics of a bacterial system with ESBL-producing and Bla-non-producing subpopulations responding to an antibiotic or a prodrug were formulated as the interactions between six main components:

$n_r$ ESBL-producing population density

$n_s$ Bla-non-producing population density

$s$ nutrient level

$a$ antibiotic concentration

$b$ extracellular ESBL concentration

$p$ prodrug concentration

$e$ released drug concentration

In this model, ESBL production extracts a fitness cost ($\alpha$) and grants resistance to the antibiotic ($\beta$) relative to the Bla-non-producing population. Nutrient is consumed by growth and released incompletely ($\xi$) with lysis. Antibiotic degradation and prodrug cleavage are mediated either by extracellular ESBLs ($\kappa_b$, $\kappa_p$) which are released on lysis, or living ESBL producers ($\varphi$, $\sigma$). All simulations were conducted in MATLAB R2023b.

$$\frac{dn_s}{d\tau} = (g - l_a)n_s \quad (1)$$

$$\frac{dn_r}{d\tau} = (\alpha g - l_m - \beta l_a)n_r \quad (2)$$

$$\frac{ds}{d\tau} = (\xi l_a - g)n_s + (\xi l_m + \xi\beta l_a - \alpha g)n_r \quad (3)$$

$$\frac{da}{d\tau} = -\kappa_b ba - \varphi n_r a - d_a a \quad (4)$$

$$\frac{db}{d\tau} = \beta l_a n_r + l_m n_r \quad (5)$$

$$\frac{dp}{d\tau} = -\kappa_p bp - \sigma n_r p \quad (6)$$

$$\frac{dm}{d\tau} = \kappa_p bp + \sigma n_r p - d_m m \quad (7)$$

$$g = \frac{s}{1+s} \tag{8}$$

$$l_a = \gamma \frac{a^{h_a}}{1+a^{h_a}} g \tag{9}$$

$$l_e = \psi \frac{m^{h_m}}{1+m^{h_m}} \tag{10}$$

The model makes several assumptions to simplify the formulation of equations: (i) Growth follows Monod growth kinetics, where nutrient level ($s$) is scaled with respect to the Monod constant. The maximum growth rate = 1/h (when nutrient is saturating and there is no burden). (ii) The antibiotic concentration ($a$) is scaled with respect to $EC_{50}$. (iii) The lysis rate from antibiotics ($l_a$) is proportional to the growth rate ($g$) with a slope that increases with the antibiotic concentration ($a$) up to a maximum lysis rate ($\gamma$). (iv) The lysis rate from the released drug ($l_m$) increases with the released drug concentration ($e$) up to a maximum lysis rate ($\psi$).

We used initial conditions of $n_s(0) = 0.2$, $n_r(0) = 0.2$, $s(0) = 4$, and $b(0) = 0$ for all simulations shown in **Fig. 1**. The antibiotic-only simulation was initialized with $a(0) = 5$ and $p(0) = 0$ and the prodrug-only simulation was initialized with $a(0) = 0$ and $p(0) = 5$. Parameter values used were as follows:

| Parameter | Significance | Value |
|---|---|---|
| $\gamma$ | antibiotic max lysis rate | 1.5 |
| $h_a$ | Hill coefficient for lysis dependence on antibiotic concentration | 3 |
| $\psi$ | released drug max lysis rate | 1.5 |
| $h_m$ | Hill coefficient for lysis dependence on released drug concentration | 2 |
| $\alpha$ | growth burden of ESBL production relative to non-producers | 0.95 |
| $\beta$ | antibiotic resistance from ESBL production relative to non-producers | 0.5 |
| $\Phi$ | antibiotic degradation rate from ESBL producers | 0.3 |
| $\sigma$ | prodrug cleavage rate from ESBL producers | 0.4 |
| $\kappa_b$ | antibiotic degradation rate from external ESBLs | 0.4 |
| $\kappa_p$ | prodrug cleavage rate from external ESBLs | 0.5 |
| $d_a$ | natural antibiotic degradation rate | 0.005 |
| $d_m$ | natural released drug degradation rate | 0.005 |
| $\xi$ | nutrient recycling inefficiency | 0.8 |

**F** **RMSDs of PBP3 Complexes from Molecular Dynamic Simulations**

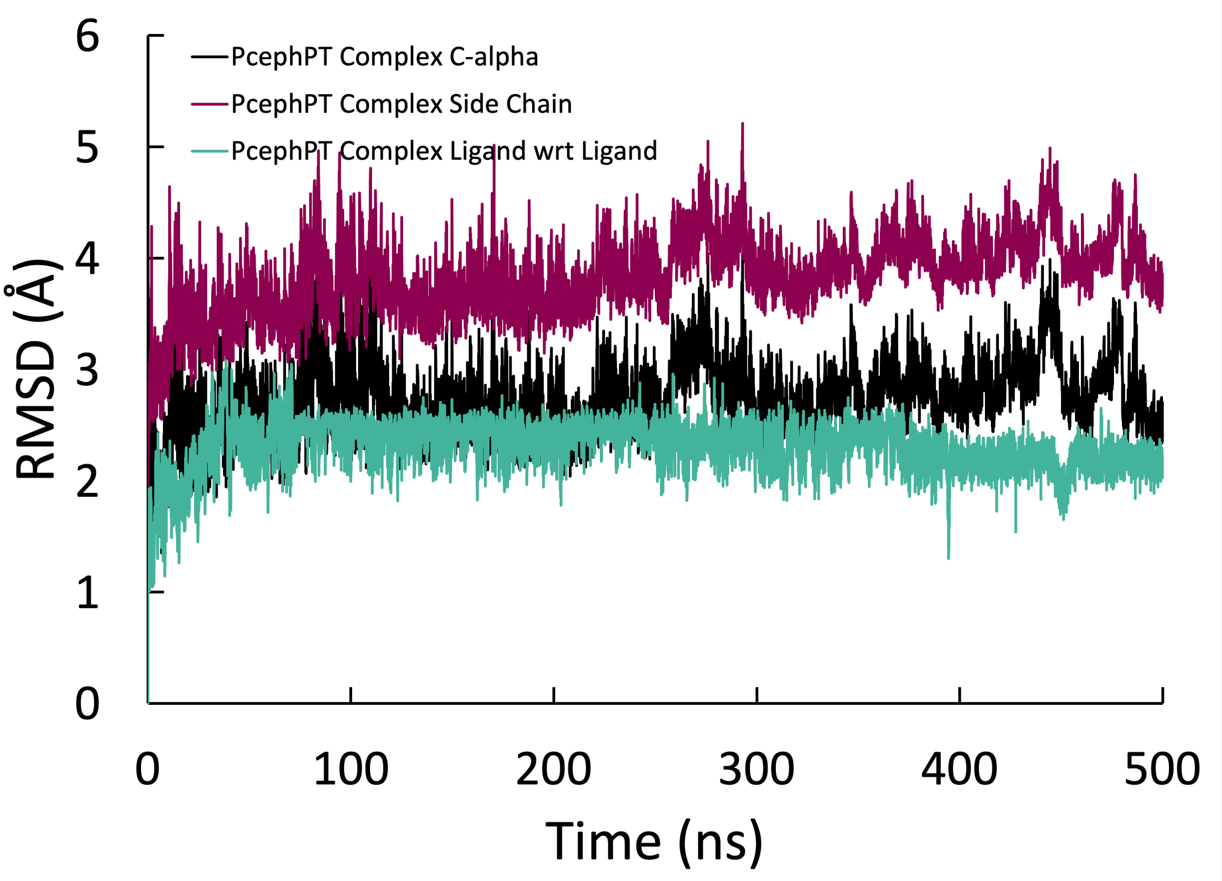


**Figure S23. RMSD plots of PcephPT/PBP3 complex confirming simulation converges within 200 ns.** RMSDs were calculated for indicated atoms of the complex using frame 1 as a reference. C-alphas is the RMSD of only the α-carbons of PBP3's backbone. Side Chains is the RMSD of only the heavy atoms of PBP3's residue side chains. Ligand with respect to (wrt) Ligand is the RMSD calculated for the heavy atoms of PcephPT in the PBP3 complex.

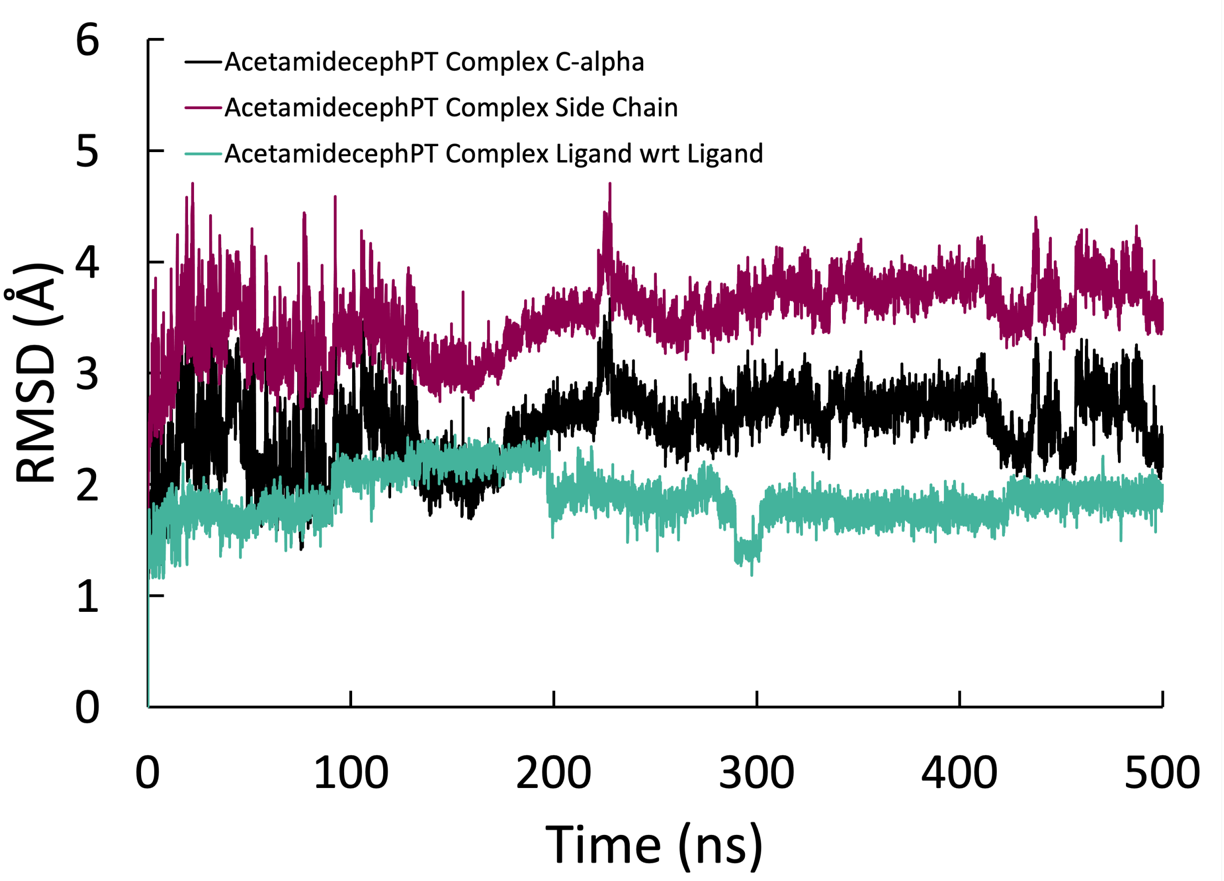


**Figure S24. RMSD plots of AcetamidecephPT/PBP3 complex confirming simulation converges within 200 ns.** RMSDs were calculated for indicated atoms of the complex using frame 1 as a reference. C-alphas is the RMSD of only the α-carbons of PBP3's backbone. Side Chains is the RMSD of only the heavy atoms of PBP3's residue side chains. Ligand with respect to (wrt) Ligand is the RMSD calculated for the heavy atoms of AcetamidecephPT in the PBP3 complex.

**G** **RMSDs of NDM-1 Complexes from Molecular Dynamic Simulations**

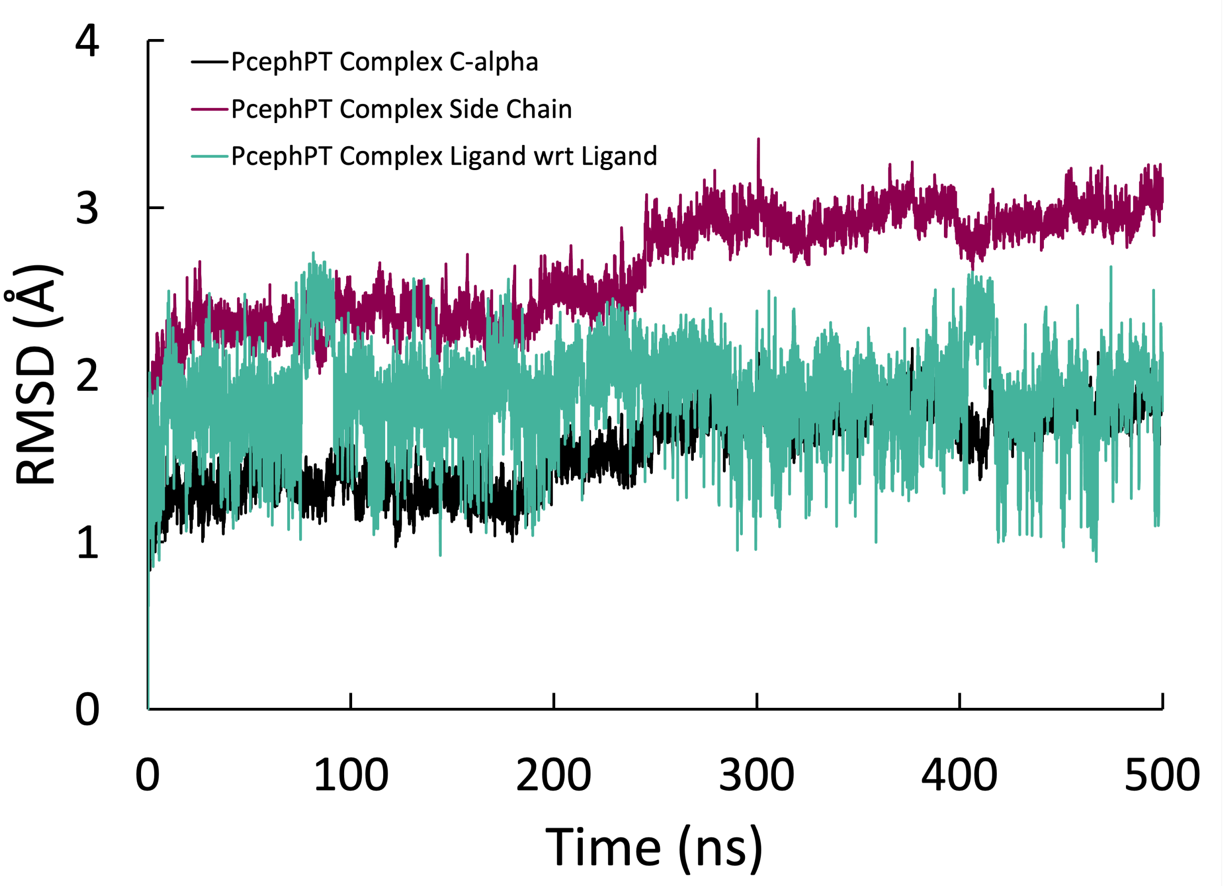


**Figure S25. RMSD plots of PcephPT/NDM-1 complex confirming simulation converges within 200 ns.** RMSDs were calculated for indicated atoms of the complex using frame 1 as a reference. C-alphas is the RMSD of only the α-carbons of NDM-1's backbone. Side Chains is the RMSD of only the heavy atoms of NDM-1's residue side chains. Ligand with respect to (wrt) Ligand is the RMSD calculated for the heavy atoms of PcephPT in the NDM-1 complex.

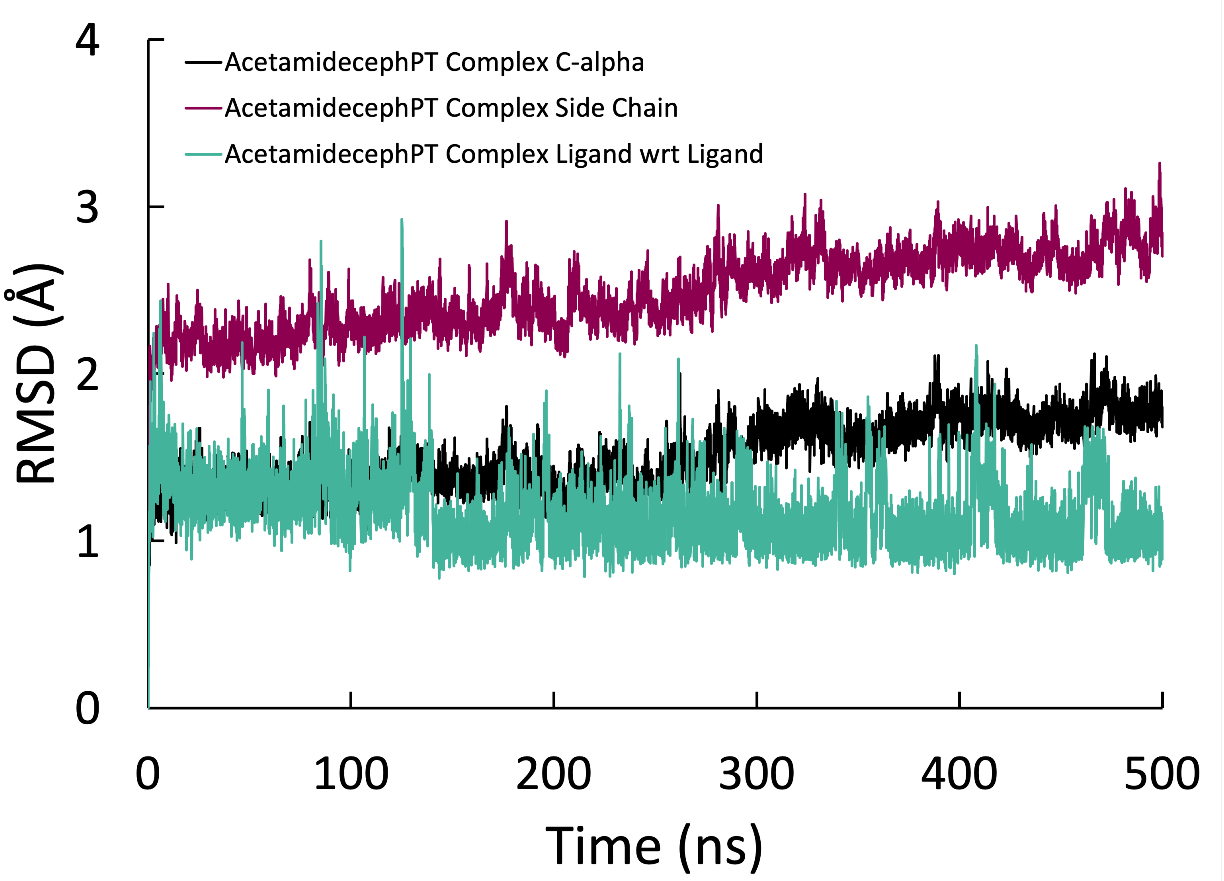


**Figure S26. RMSD plots of AcetamidecephPT/NDM-1 complex confirming simulation converges within 200 ns.** RMSDs were calculated for indicated atoms of the complex using frame 1 as a reference. C-alphas is the RMSD of only the α-carbons of NDM-1's backbone. Side Chains is the RMSD of only the heavy atoms of NDM-1's residue side chains. Ligand with respect to (wrt) Ligand is the RMSD calculated for the heavy atoms of AcetamidecephPT in the NDM-1 complex.

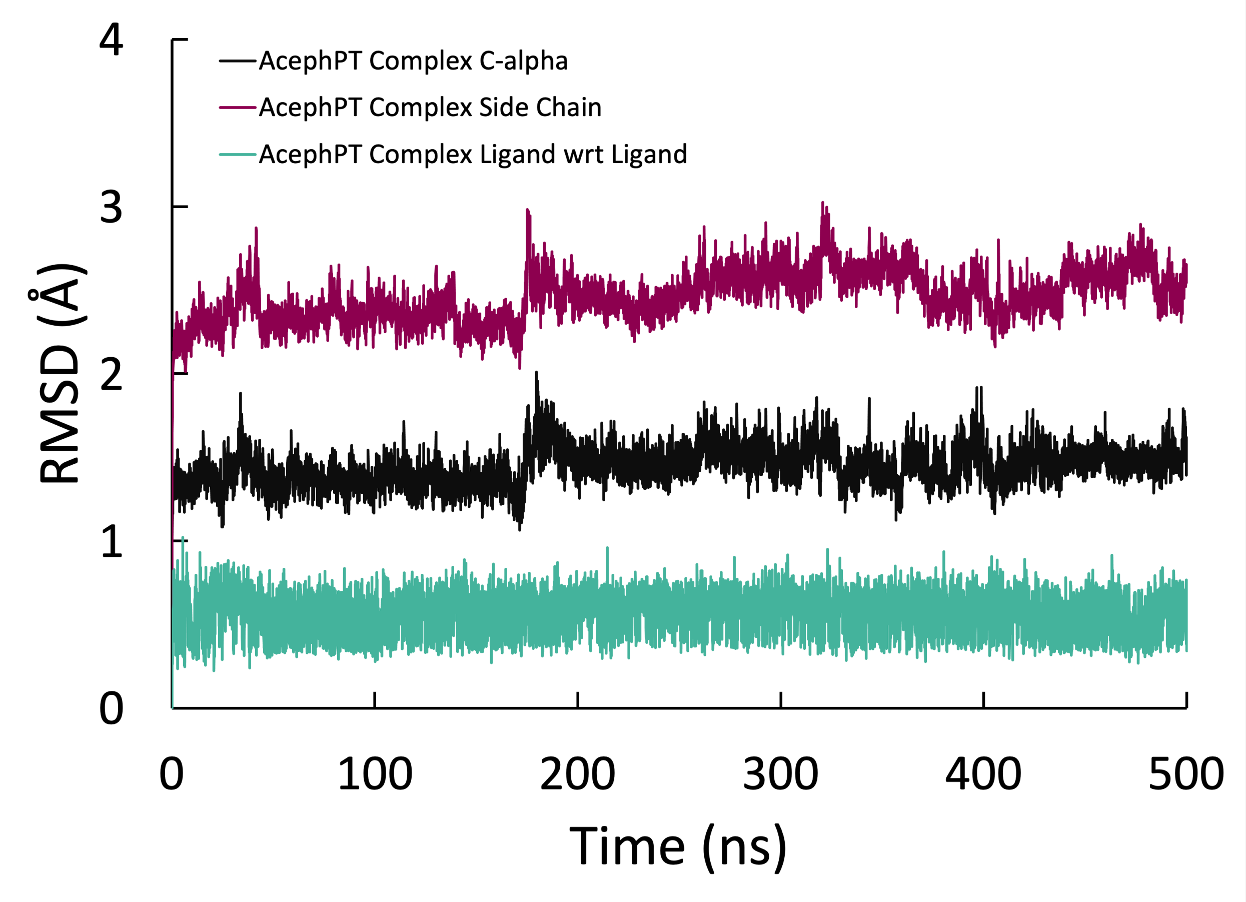


**Figure S27. RMSD plots of AcephPT/NDM-1 complex confirming simulation converges within 200 ns.** RMSDs were calculated for indicated atoms of the complex using frame 1 as a reference. C-alphas is the RMSD of only the α-carbons of NDM-1's backbone. Side Chains is the RMSD of only the heavy atoms of NDM-1's residue side chains. Ligand with respect to (wrt) Ligand is the RMSD calculated for the heavy atoms of AcephPT in the NDM-1 complex.

#### H Pyrithione Dose-Response Curves

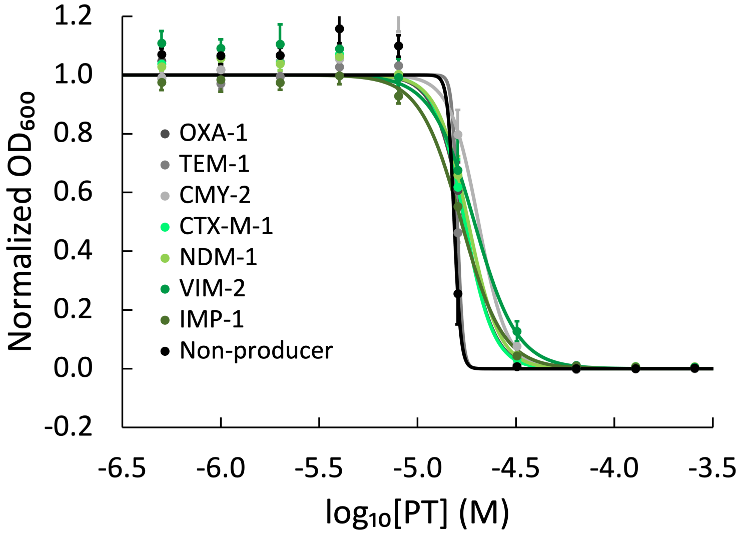


**Figure S28. Dose-response curves of pyrithione treated lab engineered *E. coli* K-12 MG1655 expressing the indicated Blas.** Blas in green are classified as ESBLs. Pyrithione suppresses the growth of all engineered strains similarly. Points and errors bars are mean ± SE from 9 biological replicates (*N* = 9 each group). Fitted lines are logistic functions determined from non-linear regression.

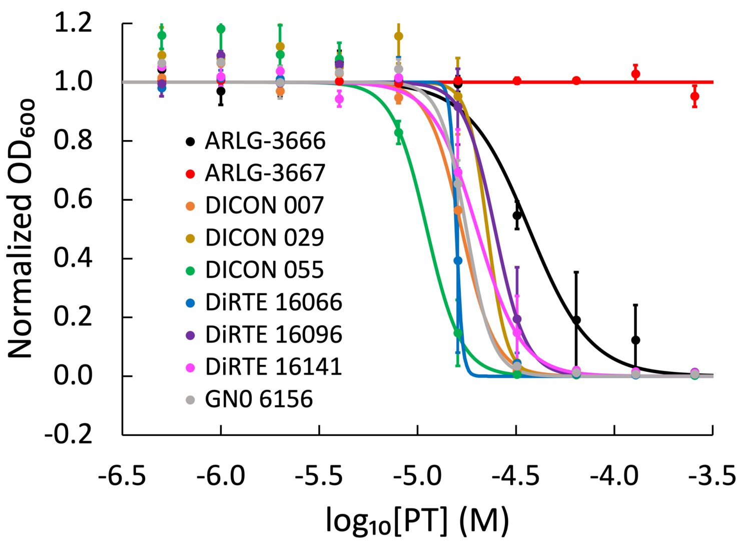


**Figure S29. Dose-response curves of pyrithione treated clincal isolates.** Points and errors bars are mean ± SE from 3 biological replicates (*N* = 3 each group). Fitted lines are logistic functions determined from non-linear regression.

#### I Clinical Isolate Characterization

**Table S1. Clinical isolate species and their expressed Blas**

| Identifier | Species | Blas Expressed | $EC_{50}$ (µM) |
|---|---|---|---|
| DiRTE 16096 | *E. coli* | NDM-1 | 29 |
| DiRTE 16141 | *E. coli* | NDM-1, OXA-48 | 46 |
| DICON 055 | *E. coli* | CTX-M-14 | 33 |
| DICON 029 | *E. coli* | CTX-M-15, OXA-1 | 267 |
| GNO 6156 | *E. coli* | CMY-2 | 84 |
| DICON 007 | *E. coli* | TEM-1 | >512 |
| DiRTE 16066 | *K. pneumoniae* | NDM-1 | 19 |
| ARLG-3666 | *E. clocae* | NDM-1, AmpC, CTX-M, TEM-1 | 74 |
| ARLG-3667 | *P. aeruginosa* | VIM-2 | >512 |

#### J Profiling the Hydrolysis of AcephPT by NDM-1-producing *E. coli*

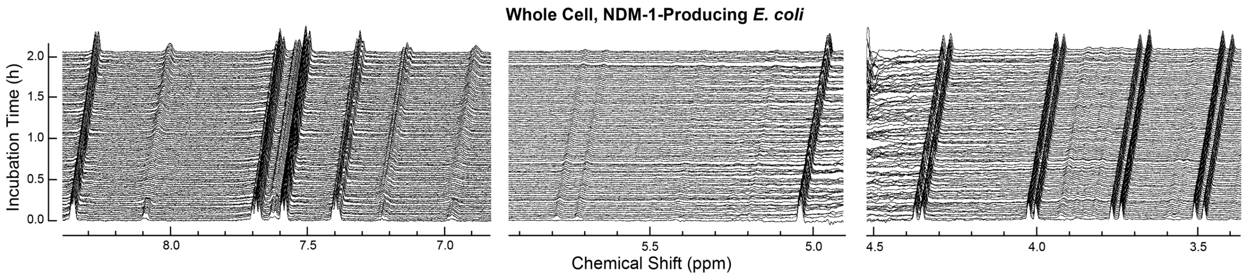


**Figure S30. Time-course NMR spectra profile the chemical changes of the prodrug AcephPT incurred upon incubation with whole cell NDM-1-producing *E. coli* at $OD_{600}$ = 2.4–2.6** NDM-1-producing *E. coli* hydrolyze an initial amount of AcephPT rapidly before the reaction stalls.

#### K Bactericidal Assays

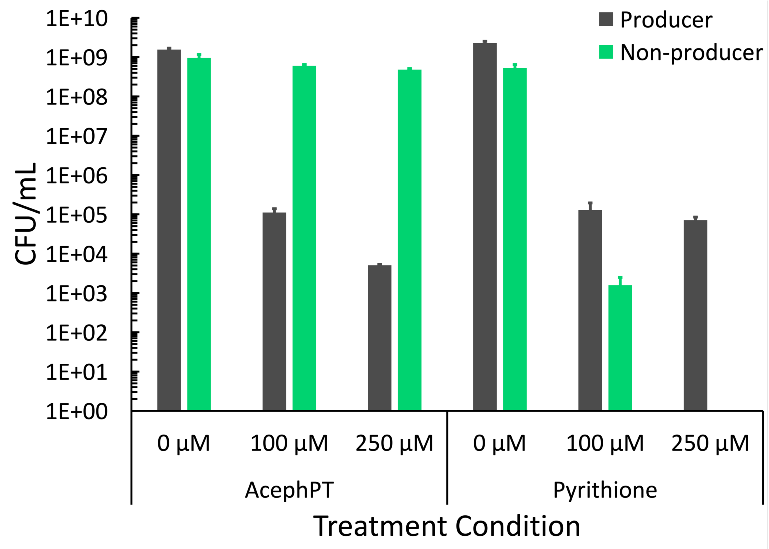


**Figure S31. Viable producing and non-producing bacteria in monoculture from treatment with AcephPT and pyrithione.** The Producer is the DiRTE 16096 clinical isolate, the Non-producer is the Top10 *E. coli* that expresses mCherry. AcephPT and pyrithione are not cytotoxic to the producer, whereas pyrithione becomes cytotoxic to the non-producer at 250 µM. Bars and error bars are mean ± SE from 3 biological replicates of 3 replicates (*N* = 9 each group).
